# A semi-supervised Bayesian approach for simultaneous protein sub-cellular localisation assignment and novelty detection

**DOI:** 10.1101/2020.05.05.078345

**Authors:** Oliver M. Crook, Aikaterini Geladaki, Daniel J.H. Nightingale, Owen Vennard, Kathryn S. Lilley, Laurent Gatto, Paul D.W. Kirk

**Affiliations:** Cambridge Centre for Proteomics, Department of Biochemistry, University of Cambridge, Cambridge, UK; MRC Biostatistics Unit, School of Clinical Medicine, University of Cambridge, Cambridge, UK; Department of Genetics, Universtiy of Cambridge, Cambridge, UK; de Duve Institute, UCLouvain, Avenue Hippocrate 75, 1200 Brussels, Belgium; Cambridge Institute of Therapeutic Immunology & Infectious Disease (CITIID), Jeffrey Cheah Biomedical Centre, Cambridge Biomedical Campus, University of Cambridge, UK

## Abstract

The cell is compartmentalised into complex micro-environments allowing an array of specialised biological processes to be carried out in synchrony. Determining a protein’s sub-cellular localisation to one or more of these compartments can therefore be a first step in determining its function. High-throughput and high-accuracy mass spectrometry-based sub-cellular proteomic methods can now shed light on the localisation of thousands of proteins at once. Machine learning algorithms are then typically employed to make protein-organelle assignments. However, these algorithms are limited by insufficient and incomplete annotation. We propose a semi-supervised Bayesian approach to novelty detection, allowing the discovery of additional, previously unannotated sub-cellular niches. Inference in our model is performed in a Bayesian framework, allowing us to quantify uncertainty in the allocation of proteins to new sub-cellular niches, as well as in the number of newly discovered compartments. We apply our approach across 10 mass spectrometry based spatial proteomic datasets, representing a diverse range of experimental protocols. Application of our approach to *hyper*LOPIT datasets validates its utility by recovering enrichment with chromatin-associated proteins without annotation and uncovers sub-nuclear compartmentalisation which was not identified in the original analysis. Moreover, using sub-cellular proteomics data from *Saccharomyces cerevisiae*, we uncover a novel group of proteins trafficking from the ER to the early Golgi apparatus. Overall, we demonstrate the potential for novelty detection to yield biologically relevant niches that are missed by current approaches.

## 1 Introduction

Aberrant protein sub-cellular localisation has been implicated in numerous diseases, including cancers (Kau *et al.*, 2004), obesity (Siljee *et al.*, 2018), and multiple others (Laurila and Vihinen, 2009). Furthermore, recent estimates suggest that up to 50% of proteins reside in multiple locations with potentially different functions in each sub-cellular niche (Christoforou *et al.*, 2016; Thul *et al.*, 2017). Characterising the sub-cellular localisation of proteins is therefore of critical importance in order to understand the pathobiological mechanisms and aetiology of many diseases. Proteins are compartmentalised into sub-cellular niches, including organelles, sub-cellular structures, liquid phase droplets and protein complexes. These compartments ensure that the biochemical conditions for proteins to function correctly are met, and that they are in the proximity of interaction partners (Gibson, 2009). A common approach to map the global sub-cellular localisation of proteins is to couple gentle cell lysis with high-accuracy mass spectrometry (MS) (Christoforou *et al.*, 2016; Mulvey *et al.*, 2017; Geladaki *et al.*, 2019; Orre *et al.*, 2019). These methods are designed to yield fractions differentially enriched in the sub-cellular compartments rather than purifying the compartments into individual fractions. As such, these spatial proteomics approaches aim to interrogate the greatest number of sub-cellular niches possible by relying upon rigorous data analysis and interpretation (Gatto *et al.*, 2010, 2014a).

Current computational approaches in MS-based spatial proteomics utilise machine learning algorithms to make protein-organelle assignments (see Gatto *et al.* (2014a) for an overview). Within this framework, novelty detection, the process of identifying differences between testing and training data, has multiple benefits. For model organisms with well annotated proteomes, novelty detection can potentially uncover groups of proteins with shared sub-cellular niches not described by the training data. Novelty detection can also prove useful in validating experimental design, either by demonstrating that contaminants have been removed or that increased resolution of organelle classes has been achieved by the experimental approach. For most non-model organisms, we have little *a priori* knowledge of their sub-cellular proteome organisation, making it challenging to curate the marker set (training dataset) from the literature (Barylyuk *et al.*, 2020). In these cases, novelty detection can assist in annotating the spatial proteome. Crucially, if a dataset is insufficiently annotated, i.e sub-cellular niches detectable in the experimental data are missing from the marker set, then this leads to the classifier making erroneous assignments, resulting in inflated *false discovery rate* (FDR) and uncertainty estimates (where available). Thus, novelty detection is a useful feature for any classifier, even if novel niche detection is not a primary aim.

Previous efforts to discover novel niches within existing sub-cellular proteomics datasets have proved valuable. Breckels *et al.* (2013) presented a phenotype discovery algorithm called *phenoDisco* to detect novel sub-cellular niches and alleviate the issue of undiscovered phenotypes. The algorithm uses an iterative procedure and the *Bayesian Injormation Criterion* (BIC) (Schwarz *et al.*, 1978) is employed to determine the number of newly detected phenotypes. Afterwards, the dataset can be re-annotated and a classifier employed to assign proteins to organelles, including those that have been newly detected. Breckels *et al.* (2013) applied their method on several datasets and discovered new organelle classes in *Arabidopsis* (Dunkley *et al.*, 2006) and *Drosophila* (Tan *et al.*, 2009). This approach later successfully identified the trans-Golgi network (TGN) in *Arabidopsis* roots (Groen *et al.*, 2014).

Recent work has demonstrated the importance of uncertainty quantification in spatial proteomics (Crook *et al.*, 2018, 2019a,b). Crook *et al.* (2018) proposed a generative classification model and took a Bayesian approach to spatial proteomics data analysis by computing probability distributions of protein-organelle assignments using Markov-chain Monte-Carlo (MCMC). These probabilities were then used as the basis for organelle allocations, as well as to quantify the uncertainty in these allocations. On the basis that some proteins cannot be well described by any of the annotated sub-cellular niches, a multivariate Student’s T distribution was included in the model to enable outlier detection. The proposed T-Augmented Gaussian Mixture (TAGM) model was shown to achieve state-of-the-art predictive performance against other commonly used machine learning algorithms (Crook *et al.*, 2018). Furthermore, the model has been successfully applied to reveal unrivalled insight into the spatial organisation of *Toxoplasma gondii* (Barylyuk *et al.*, 2020) and identify cargo of the Golgins of the trans-Golgi network (Shin *et al.*, 2019).

Here, we propose an extension to TAGM to allow simultaneous protein-organelle assignments and novelty detection. One assumption of the existing TAGM model is that the number of sub-cellular niches is known. Here, we design a novelty detection algorithm based on allowing an unknown number of additional sub-cellular niches, as well as quantifying uncertainty in this number.

Quantifying uncertainty in the number of clusters in a Bayesian mixture model is challenging and many approaches have been proposed in the literature (see for example Ferguson (1974); Antoniak (1974); Richardson and Green (1997) and the appendix for further details). Here, we make use of asymptotic results in Bayesian analysis of mixture models (Rousseau and Mengersen, 2011). The principle of *overfitted mixtures* allows us to specify a (possibly large) maximum number of clusters. As shown in Rousseau and Mengersen (2011) these components empty if they are not supported by the data, allowing the number of clusters to be inferred. Kirk *et al.* (2012) previously made use of this approach in the Bayesian integrative modelling of multiple genomic datasets. In our application, some of the organelles may be annotated with known marker proteins and this places a lower bound on the number of sub-cellular niches. Bringing these ideas together results in a semi-supervised Bayesian approach, which we refer to as Novelty TAGM. Table 1 summarises the differences between the current available machine-learning methods for spatial proteomics.

**Table 1:** Examples of computational methods for spatial proteomics datasets for prediction and novelty detection.

| MS-based Spatial Proteomics Computational Methods for Prediction and Novelty Detection |  |  |  |  |  |  |  |
| --- | --- | --- | --- | --- | --- | --- | --- |
| Method | Localisation prediction | Uncertainty in protein localisation | Outlier detection | Novelty detection | Uncertainty in number of novel phenotypes | Uncertainty in allocation to new phenotypes | Integrative |
| Supervised Machine Learning (as reviewed in Gatto et al. (2014a)) | ✓ | ✗ | ✗ | ✗ | ✗ | ✗ | ✗ |
| Correlation Profiling (Foster et al., 2006; Krahmer et al., 2018) | ✓ | ✗ | ✗ | ✗ | ✗ | ✗ | ✗ |
| Transfer Learning (Breckels et al., 2016) | ✓ | ✗ | ✗ | ✗ | ✗ | ✗ | ✓ |
| *Mclust* (as used in Orre et al. (2019)) | ✗ | ✗ | ✓ | ✓ | ✗ | ✗ | ✗ |
| *PhenoDisco* (Breckels et al., 2013) | ✗ | ✗ | ✓ | ✓ | ✗ | ✗ | ✗ |
| TAGM (Crook et al., 2018) | ✓ | ✓ | ✓ | ✗ | ✗ | ✗ | ✗ |
| Novelty TAGM (This manuscript) | ✓ | ✓ | ✓ | ✓ | ✓ | ✓ | ✗ |

We apply Novelty TAGM to 10 spatial proteomic datasets across a diverse range of protocols, including *hyper*LOPIT (Christoforou *et al.*, 2016; Mulvey *et al.*, 2017), LOPIT-DC (Geladaki *et al.*, 2019), Dynamic Organellar Maps (DOM) (Itzhak *et al.*, 2016) and spatial-temporal methods (Beltran *et al.*, 2016). Application of Novelty TAGM to each dataset reveals additional biologically relevant compartments. Notably, we detect 4 sub-nuclear compartments in the the U-2 OS *hyper*LOPIT dataset: the nucleolus, nucleoplasm, chromatin-associated, and the nuclear membrane. In addition, an endosomal compartment is robustly identified across *hyper* LOPIT and LOPIT-DC datasets. Finally, we also uncover collections of proteins with previously uncharacterised localisation patterns; for example, vesicle proteins trafficking from the ER to the early Golgi in *Saccharomyces cerevisiae*.

## 2 Methods

### 2.1 Datasets

We provide a brief description of the datasets used in this manuscript. We analyse *hyper*LOPIT data, in which sub-cellular fractionation is performed using density-gradient centrifugation (Dunkley *et al.*, 2004, 2006; Mulvey *et al.*, 2017), on pluripotent mESCs (E14TG2a) (Christoforou *et al.*, 2016), human bone osteosarcoma (U-2 OS) cells (Thul *et al.*, 2017; Geladaki *et al.*, 2019), and *S. cerevisiae* (bakers’ yeast) cells (Nightingale *et al.*, 2019). The mESC dataset combines two 10-plex biological replicates and quantitative information on 5032 proteins. The U-2 OS dataset combines three 20-plex biological replicates and provides information on 4883 proteins. The yeast dataset represents four 10-plex biological replicate experiments performed on *S. cerevisiae* cultured to early-mid exponential phase. This dataset contains quantitative information for 2846 proteins that were common across all replicates. Tandem Mass Tag (TMT) (Thompson *et al.*, 2003) labelling was used in all *hyper*LOPIT experiments with LC-SPS-MS^3^ used for high accuracy quantitation (Ting *et al.*, 2011; McAlister *et al.*, 2014). Beltran *et al.* (2016) integrated a temporal component to the LOPIT protocol. They analysed HCMV-infected primary fibroblast cells over 5 days, producing control and infected maps every 24 hours. We analyse the control and infected maps 24 hours post-infection, providing information on 2220 and 2196 proteins respectively. In a comparison with *phenoDisco*, we apply Novelty TAGM to a dataset acquired using LOPIT-based fractionation and 8-plex iTRAQ labelling on the HEK-293 human embryonic kidney cell line, quantifying 1371 proteins (Breckels *et al.*, 2013).

Our approach is not limited to spatial proteomics data where the sub-cellular fractionation is performed using density gradients. We demonstrate this through the analysis of DOM datasets on HeLa cells and mouse primary neurons (Itzhak *et al.*, 2016, 2017), which quantify 3766 and 8985 proteins respectively. These approaches used SILAC quantitation with differential centrifugation-based fractionation. We analyse 6 replicates from the HeLa cell line analyses in Itzhak *et al.* (2016) and 3 replicates from the mouse primary neuron experiments in Itzhak *et al.* (2017). Hirst *et al.* (2018) also used the DOM protocol coupled with CRISPR-CAS9 knockouts in order to explore the functional role of AP-3. We analyse the control map from this experiment. Finally, we consider the U-2 OS data which were acquired using the LOPIT-DC protocol (Geladaki *et al.*, 2019) and quantified 6837 proteins across 3 biological replicates. In favour of brevity, we do not consider protein correlation profiling (PCP) based spatial proteomics datasets in this manuscript, though our method also applies to such data (Foster *et al.*, 2006; Kristensen *et al.*, 2012; Kristensen and Foster, 2014) and other sub-cellular proteomics methods which utilised cellular fractionation (Orre *et al.*, 2019).

### 2.2 Model

#### 2.2.1 Spatial proteomics mixture model

In this section, we briefly review the TAGM model proposed by Crook *et al.* (2018). Let *N* denote the number of observed protein profiles each of length *L*, corresponding to the number of quantified fractions. The quantitative profile for the *i*-th protein is denoted by **x**_*i*_ = [*x*_1*i*_, …, *x_Li_*]. In the original formulation of the model it is supposed that there are *K* known sub-cellular compartments to which each protein could be localised (e.g. cytosol, endoplasmic reticulum, mitochondria, …). For simplicity of exposition, we refer to these *K* sub-cellular compartments as *components*, and introduce component labels *z_i_*, so that *z_i_* = *k* if the *i*-th protein localises to the *k*-th component. To fix notation, we denote by *X_L_* the set of proteins whose component labels are known, and by *X_U_* the set of unlabelled proteins. If protein *i* is in *X_U_*, we seek to evaluate the probability that *z_i_* = *k* for each *k* = 1, …, *K*; that is, for each unlabelled protein, we seek the probability of belonging to each component (given a model and the observed data).

The distribution of quantitative profiles associated with each protein that localises to the *k*-th component is modelled as multivariate normal with mean vector ***μ***_*k*_ and covariance matrix Σ_*k*_. However, many proteins are dispersed and do not fit this assumption. To model these “outliers”, Crook *et al.* (2018) introduced a further indicator variable *ϕ*. Each protein **x**_*i*_ is then described by an additional variable *ϕ_i_*, with *ϕ_i_* = 1 indicating that protein **x**_*i*_ belongs to an organelle-derived component and *ϕ_i_* = 0 indicating that protein **x**_*i*_ is not well described by these known components. This *outlier component* is then modelled as a multivariate T distribution with degrees of freedom *κ*, mean vector **M**, and scale matrix *V*. Thus the model can be written as:

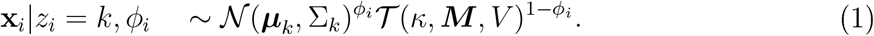

Let *f* (**x**|***μ***, Σ) denote the density of the multivariate normal with mean vector ***μ*** and covariance matrix Σ evaluated at **x**, and similarly let *g*(**x**|*κ,* **M**, **V**) denote the density of the multivariate T-distribution. For any *i*, the prior probability of the *i*-th protein localising to the *k*-th component is denoted by *p*(*z_i_* = *k*) = *π_k_*. Letting 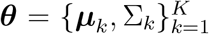 denote the set of all component mean and covariance parameters, and 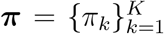 denote the set of all mixture weights, it follows that:

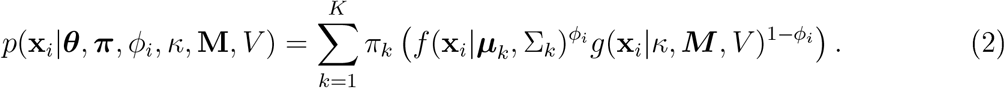

For any *i*, we set the prior probability of the *i*-th protein belonging to the outlier component as *p*(*ϕ_i_* = 0) = *ϵ*, where *ϵ* is a parameter that we infer.

Equation (2) can then be rewritten in the following way:

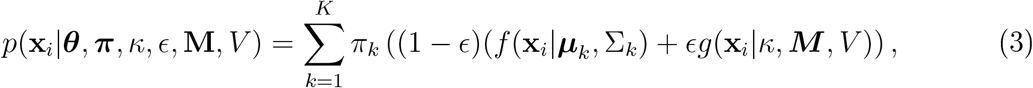

As in Crook *et al.* (2018), we fix *κ* = 4, **M** as the global empirical mean, and *V* as half the global empirical variance of the data, including labelled and unlabelled proteins. To extend this model to permit novelty detection, we specify the maximum number of components *K_max_ > K*. Our proposed model then allows up to *K_novelty_* = *K_max_ — K* ≥ 0, new phenotypes to be detected. Equation 3 can then be written as

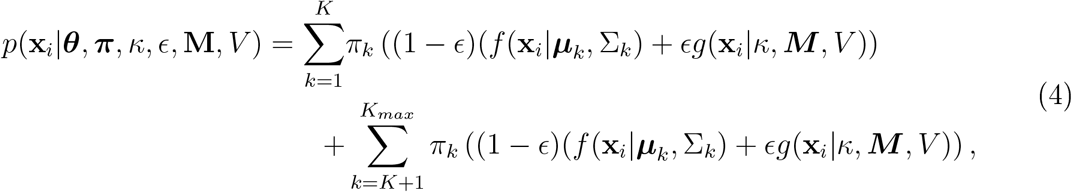

where, in the first summation, the *K* components correspond to known sub-cellular niches and the second summation corresponds to the new phenotypes to be inferred. The parameter sets are then augmented to include these possibly new components; that is, we redefine 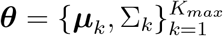 to denote the set of all component mean and covariance parameters, and 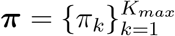 denotes the set of all mixture weights. Relying on the principle of over-fitted mixtures (Rousseau and Mengersen, 2011), components that are not supported by the data are left empty with no proteins allocated to them. We find setting *K_novety_* = 10 is ample to detect new phenotypes. To complete our Bayesian model, we need to specify priors. Detailed prior specifications and sensitivity analysis are provided in the supplement (section 7.9).

#### 2.2.2 Bayesian inference and convergence

Ve perform Bayesian inference using Markov-chain Monte-Carlo methods. We make modifications to the collapsed Gibbs sampler approach used previously in Crook *et al.* (2018) to allow inference to be performed for the parameters of the novel components (see supplement for full details). Since the number of occupied components at each iteration is random, we monitor this quantity as a convergence diagnostic.

#### 2.2.3 Visualising patterns in uncertainty

To simultaneously visualise the uncertainty in the number of newly discovered phenotypes, as well as the uncertainty in the allocation of proteins to new components, we use the so-called *posterior similarity matrix* (PSM) (Fritsch and Ickstadt, 2009). The PSM is an *N × N* matrix where the (*i, j*)^*th*^ entry is the posterior probability that protein *i* and protein *j* reside in the same component. Throughout we use a heatmap representation of this quantity. The PSM is summarised into a clustering by maximising the posterior expected adjusted Rand index (see appendix for details; (Fritsch and Ickstadt, 2009)). Formulating inference around the PSM also avoids some technical statistical challenges, which are discussed in detail in the appendix.

#### 2.2.4 Uncertainty quantification

We may be interested in quantifying the uncertainty in whether a protein belongs to a new sub-cellular component. Indeed, it is important to distinguish whether a protein belongs to a new phenotype or if we simply have large uncertainty about its localisation. The probability that protein *i* belongs to a new component is computed from the following equation:

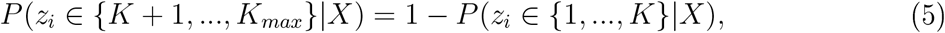

which we approximate by the following Monte-Carlo average:

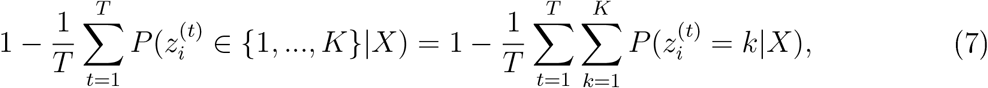

where *T* is the number of Monte-Carlo iterations. Throughout, we refer to equation 7 as the *discovery probability*.

#### 2.2.5 Applying the model in practice

Applying Novelty TAGM to spatial proteomics datasets consists of several steps. After having run the algorithm on a dataset and assessing convergence, we proceed to explore the ouput of the method. We explore *putative phenotypes*, which we define as newly discovered clusters with at least 1 protein with discovery probability greater than 0.95.

### 2.3 Validating computational approaches

In a supervised framework the performance of computational methods can be assessed by using the training data, where a proportion of the training data is withheld from the classifier to be used for the assessment of predictive performance. In an unsupervised or semi-supervised framework we cannot validate in this way, since there is no “ground truth” with which to compare. Thus, we propose several approaches, using external information, for validation of our method.

#### 2.3.1 Artificial masking of annotations to recover experimental design

Removing the labels from an entire component and assessing the ability of our method to rediscover these labels is one form of validation. We consider this approach for several of the datasets; in particular, chromatin enrichment was performed in two of the *hyper*LOPIT experiments, where the intention was to increase the resolution between chromatin and non-chromatin associated nuclear proteins (Christoforou *et al.*, 2016; Mulvey *et al.*, 2017; Thul *et al.*, 2017). As validation of our method we hide these labels and seek to rediscover them in an unbiased fashion.

#### 2.3.2 The Human Protein Atlas

A further approach to validating our method is to use additional spatial proteomic information. The Human Protein Atlas (HPA) (Thul *et al.*, 2017; Sullivan *et al.*, 2018) provides confocal microscopy information on thousands of proteins, using validated antibodies. When we consider a dataset for which there is HPA annotation, we use this data to validate the novel phenotypes for biological relevance.

#### 2.3.3 Gene Ontology (GO) term enrichment

Throughout, we perform GO enrichment analysis with FDR control performed according to the Benjamini-Höchberg procedure (Benjamini and Hochberg, 1993; Ashburner *et al.*, 2000; Yu *et al.*, 2012). The proteins in each novel putative phenotype are assessed in turn for enriched Cellular Component terms, against the background of all quantified proteins in that experiment.

#### 2.3.4 Robustness across multiple MS-based spatial proteomics datasets

On occasion some cell lines have been analysed using multiple spatial proteomics technologies (Geladaki *et al.*, 2019). In these cases, the putative phenotypes discovered by Novelty TAGM are compared directly. If the same phenotype is discovered in different proteomic datasets we consider this as robust evidence for sufficient resolution of that phenotype.

## 3 Results

Motivated by the need for novelty detection methods which also quantify the uncertainty in the number of clusters and the assignments of proteins to each cluster, we developed Novelty TAGM (see methods). This approach extends our previous TAGM model (Crook *et al.*, 2018) to enable the detection of novel *putative phenotypes*, which we define as newly discovered clusters with at least 1 protein with discovery probability greater than 0.95. Our proposed methodology allows us to interrogate individual proteins to assess whether they belong to a newly discovered phenotype. Through the *posterior similarity matrix* (PSM) we can visualise the global patterns in the uncertainty in phenotype discovery (see supplement). We summarise this posterior similarity matrix into a single clustering by maximising the posterior expected adjusted rand index (see methods). This methodology infers the number of clusters supported by the data, in contrast to many existing approaches which require specification of the number of clusters (such as K-means or Mclust (Fraley *et al.*, 2012)). To demonstrate the value of this approach, we applied Novelty TAGM to a diverse set of spatial proteomics datasets.

### 3.1 Validating experimental design in *hyper*LOPIT

Initially, we validated Novelty TAGM in a setting where we have a strong *a priori* expectation for the presence of an unannotated niche. For this we used a human bone osteosarcoma cell (U-2 OS) *hyper*LOPIT dataset (Thul *et al.*, 2017) and an mESC *hyper*LOPIT dataset (Christoforou *et al.*, 2016). These experimental protocols used a chromatin enrichment step to resolve nuclear chromatin-associated proteins from nuclear proteins not associated with chromatin. Removing the nuclear, chromatin and ribosomal annotations from the datasets, we test the ability of Novelty TAGM to recover them.

#### 3.1.1 Human bone osteosarcoma (U-2 OS) cells

For the U-2 OS dataset, Novelty TAGM reveals 9 putative phenotypes, which we refer to as phenotype 1, phenotype 2, etc… These phenotypes, along with the uncertainty associated with them, are visualised in figure 2. We consider the HPA confocal microscopy data for validation (Thul *et al.*, 2017; Sullivan *et al.*, 2018). The HPA provides information on the same cell line and therefore constitutes an excellent complementary resource. This *hyper*LOPIT dataset was already shown to be in strong agreement with the microscopy data (Thul *et al.*, 2017; Geladaki *et al.*, 2019). Proteins in phenotypes 3, 4, 5 and 8 have a nucleus-related annotation as their most frequent HPA annotation, as well as differential enrichment of nucleus-related GO terms (figure 2). Phenotype 3 validates the chromatin enrichment preparation (figure 2 panel (c)) and phenotype 4 reveals a nucleoli cluster, where nucleoli and nucleoli/nucleus are the 2^*nd*^ and 3^*rd*^ most frequent HPA annotations for proteins belonging to this phenotype. For phenotype 5, the most associated term is nucleoplasm from the HPA data and this is further supported by GO analysis (figure 2 panel (c)). Phenotype 8 demonstrates further sub-nuclear resolution and has *nuclear membrane* as its most frequent HPA annotation and has corresponding enriched GO terms (figure 2 panel (c)). In addition, phenotypes 1 and 2 are enriched for *ribosomes* and *endosomes* respectively.

**Figure 1:**
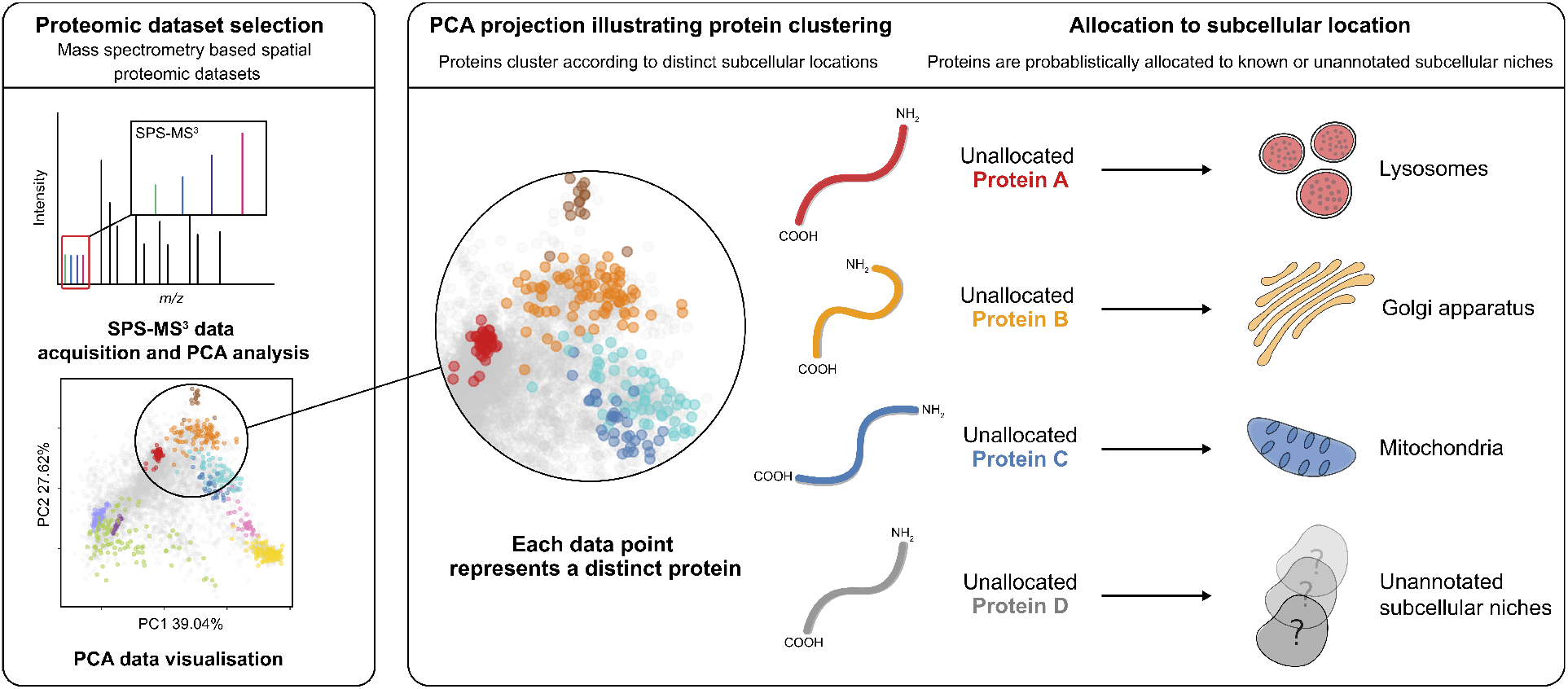
An overview of novelty detection in subcellular proteomics.

**Figure 2:**
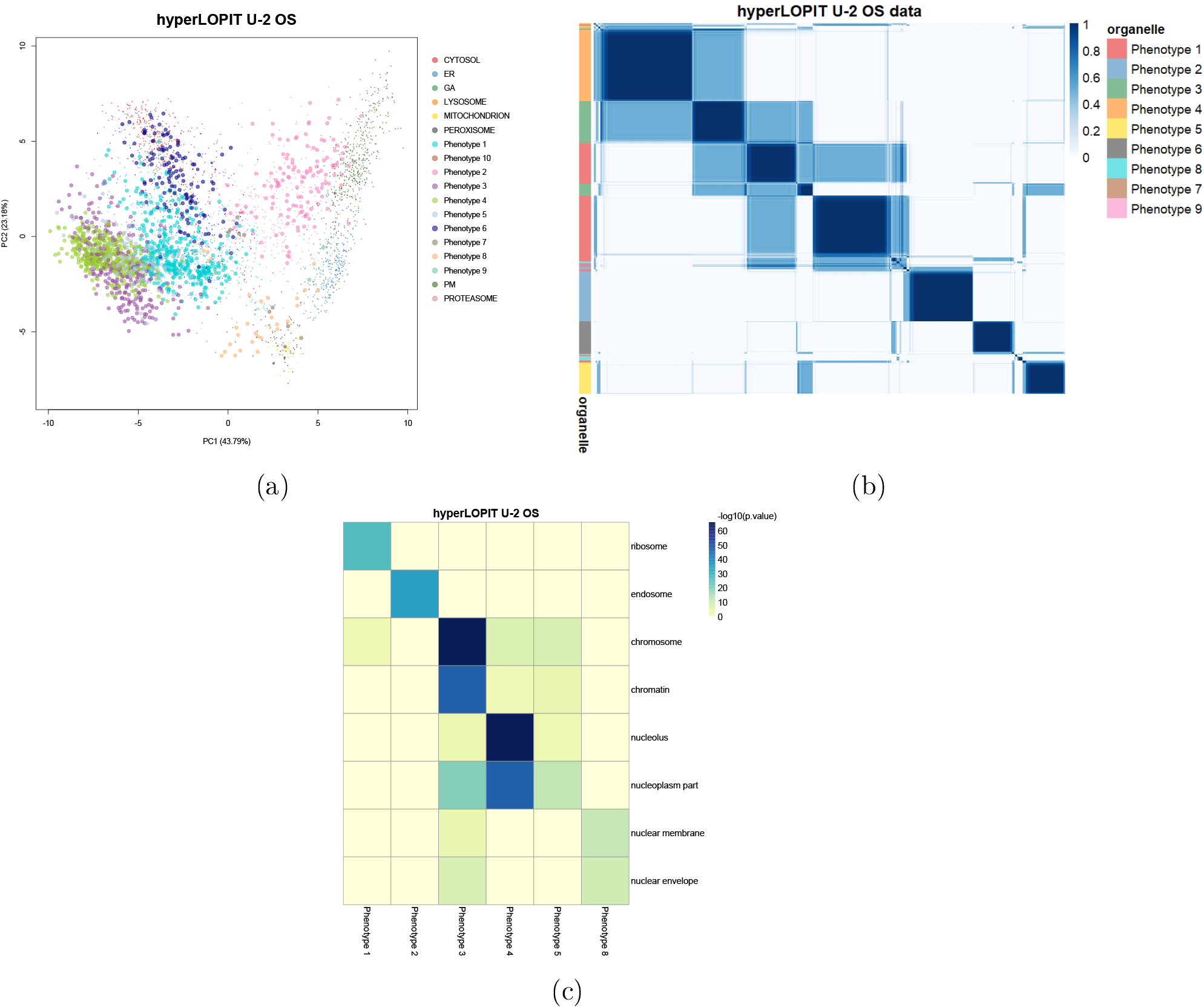
(a) PCA plot of the *hyper*LOPIT U-2 OS cancer cell line data. Points are scaled according to the discovery probability with larger points indicating greater discovery probability. (b) Heatmaps of the posterior similarity matrix derived from U-2 OS cell line data demonstrating the uncertainty in the clustering structure of the data. We have only plotted the proteins which have greater than 0.99 probability of belonging to a new phenotype and probability of being an outlier less than 0.5 for the U-2 OS dataset to reduce the number of visualised proteins. (c) Tile plot of discovered phenotypes against GO CC terms to demonstrate over-representation, where the colour intensity is the −log_10_ of the p-value.

#### 3.1.2 Pluripotent mESCs (E14TG2a)

In the case of the mESC dataset, Novelty TAGM reveals 8 new putative phenotypes. The chromatin enrichment preparation is also validated in these cells, as well as new phenotypes with additional annotations such *nucleolus* and *centrosome* (see supplement 7.1). We also used this dataset to explore how our results are impacted if we reduce the number of markers from other niches (see supplement 7.10).

### 3.2 Uncovering additional sub-cellular structures

Having validated the ability of Novelty TAGM to recover known experimental design, as well as uncover additional sub-cellular niches resolved in the data, we turn to apply Novelty TAGM to several additional datasets.

#### 3.2.1 U-2 OS cell line revisited

We first consider the LOPIT-DC dataset on the U-2 OS cell line (Geladaki *et al.*, 2019). Again, we removed the nuclear, proteasomal, and ribosomal annotations. Novelty TAGM reveals 10 putative phenotypes (figure 3).

**Figure 3:**
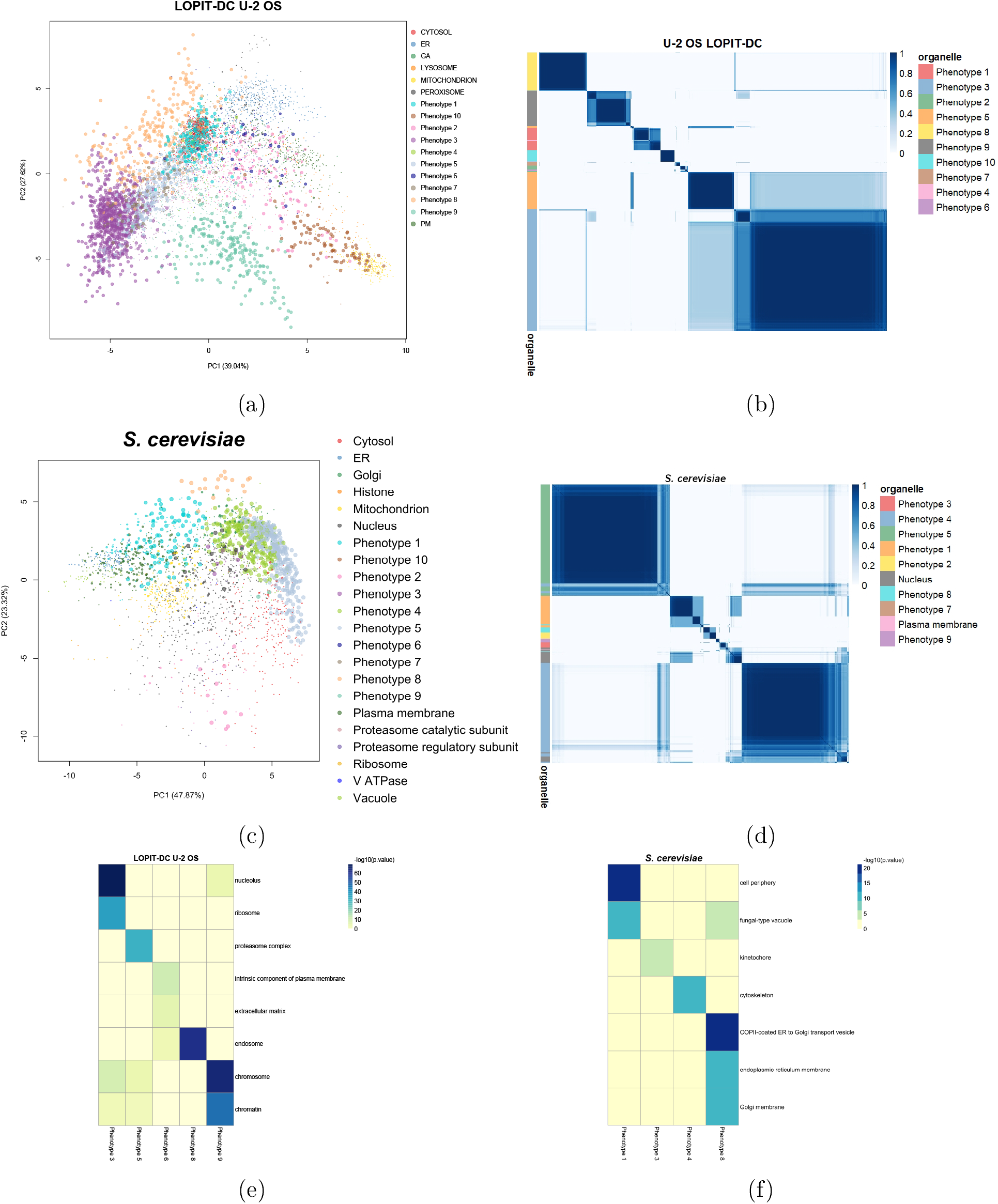
(a, c) PCA plots of the LOPIT-DC U-2 OS data and the *hyper*LOPIT yeast data. The points are scaled according to the discovery probability. (b, d) Heatmaps of the posterior similarity matrix derived from the U-2 OS and yeast datasets demonstrating the uncertainty in the clustering structure of the data. We have only plotted the proteins which have greater than 0.99 probability of belonging to a new phenotype and probability of being an outlier less than 0.93 (10^−5^ for LOPIT-DC to reduce the number of visualised proteins). (e, f) Tile plots of phenotypes against GO CC terms where the colour intensity is the −log_10_ of the p-value.

In a similar vein to the analysis performed on the *hyper*LOPIT U-2 OS dataset, we initially use the available HPA data to validate these clusters (Thul *et al.*, 2017). Phenotypes 3, 5, 7 and 9 display nucleus-associated terms as their most frequent HPA annotation. Clear differential enrichment of phenotypes with GO Cellular Component terms is evident from figure 3 panel (e). This analysis reveals *nucleolus, ribosome, proteasome* phenotypes. Furthermore, a *chromatin* phenotype is also resolved. Notably, this is the first evidence for sub-nuclear resolution in this LOPIT-DC dataset. Phenotype 6 represents a cluster with mixed *plasma membrane* and *extracellular matrix* annotations and this is supported by HPA annotation with vesicles, cytosol, and plasma membrane being the top three annotations. An extracellular matrix-related phenotype was not previously known in these data and might correspond to exocytic vesicles containing ECM proteins. Furthermore, phenotype 8 is significantly enriched for *endosomes*, again a novel annotation for this data. In addition, 107 of the proteins in this phenotype are also localised to the endosome-enriched phenotype presented in the U-2 OS *hyper*LOPIT dataset (section 3.1.1). Thus, we robustly identify new phenotypes across different spatial proteomics protocols. Hence, we have presented strong evidence for additional annotations in this dataset, beyond the original analysis of the data (Geladaki *et al.*, 2019). In particular, although a separate chromatin enrichment preparation was not included in the U-2 OS LOPIT-DC analysis and the original authors did not identify sufficient resolution between the nucleus and chromatin clusters in this dataset, Novelty TAGM could, in fact, reveal a chromatin-associated phenotype in the U-2 OS LOPIT-DC data. In addition, we have joint evidence for an endosomal cluster in both the LOPIT-DC and *hyper*LOPIT datasets. Finally, through the discovery probability and by using the PSMs we have quantified uncertainty in these proposed phenotypes, enabling more rigorous interrogation of these datasets.

#### 3.2.2 Saccharomyces cerevisiae

Novelty TAGM uncovers 8 putative phenotypes in the yeast *hyper*LOPIT data (Nightingale *et al.*, 2019). Four of these phenotypes have no significant over-represented annotations. Figure 3 panel (f) demonstrates that the remaining four phenotypes are differentially enriched for GO terms. Firstly, a mixed *cell periphery* and *fungal-type vacuole* phenotype is uncovered along with a *kinetochore* phenotype, and a *cytoskeleton* phenotype. Phenotype 8 represents a joint Golgi and ER cluster with several enriched GO terms. Indeed, most of the proteins in this phenotype have roles in the early secretory pathway that involve either transport from the ER to the early Golgi apparatus, or retrograde transport from the Golgi to the ER (Bue *et al.*, 2006; Inadome *et al.*, 2005; Otte *et al.*, 2001; Yofe *et al.*, 2016), (also reviewed in Delic *et al.* (2013)). To be precise, 11 out of the total 20 proteins in this cluster are annotated as core components of COPII vesicles and 6 associated with COPI vesicles. The protein Ksh1p (Q8TGJ3) is further suggested through homology with higher organisms to be part of the early secretory pathway (Wendler *et al.*, 2010). The proteins Scw4p (P33334), Cts1p (P29029) and Scw10p (Q04931) (Cappellaro *et al.*, 1998), as well as Pst1p (Q12333)(Pardo *et al.*, 2004), and Cwp1p (P28319) (Yin *et al.*, 2005), however, are annotated in the literature as localising to the cell wall or extracellular region. It is therefore possible that their predicted co-localisation with secretory pathway proteins observed here reflects a proportion of their lifecycle being synthesised or spent trafficking through the secretory pathway. The protein Ssp120p (P39931) is of unknown function and has been shown to localise in high throughput studies to the vacuole (Yofe *et al.*, 2016) and to the cytoplasm in a punctate pattern (Huh *et al.*, 2003). The localisation observed here may suggest that it is therefore either part of the secretory pathway, or trafficks through the secretory organelles for secretion or to become a constituent of the cell wall.

#### 3.2.3 Fibroblast cells

We also uncover additional annotations for the HCMV infected and the control fibroblast spatial proteomics datasets (Beltran *et al.*, 2016); such as, sub-mitochondrial annotations, as well as resolution of the small and large ribosomal sub-units. These annotations were overlooked in the original analysis (Beltran *et al.*, 2016) and further details can be found in the supplement 7.2.

### 3.3 Refining annotation in organellar maps

The Dynamic Organellar Maps (DOM) protocol was developed as a faster method for MS-based spatial proteomic mapping, albeit at the cost of lower organelle resolution (Itzhak *et al.*, 2016; Gatto *et al.*, 2019). The three datasets analysed here are two HeLa cell lines (Itzhak *et al.*, 2016; Hirst *et al.*, 2018) and a mouse primary neuron dataset (Itzhak *et al.*, 2017). All three of these datasets have been annotated with a class called “large protein complexest”. This class contains a mixture of cytosolic, ribosomal, proteasomal and nuclear sub-compartments that pellet during the centrifugation step used to capture this mixed fraction (Itzhak *et al.*, 2016). We apply Novelty TAGM to these data and remove this “large protein complexes” class, to derive more precise annotations for these datasets.

#### 3.3.1 HeLa cells (Itzhak et. al 2016)

The HeLa dataset of Itzhak *et al.* (2016) has 3 additional phenotypes uncovered by Novelty TAGM. Figure 4 panel c shows a *mitochondrial membrane* phenotype, distinct from the already annotated mitochondrial class. Phenotype 2 represents a mixed cluster with nucleus-, ribosome- and cytosol-related enriched terms. The final phenotype is enriched for *chromatin* and *chromosome*, suggesting sub-nuclear resolution. Furthermore, as a result of quantifying uncertainty, we can see that there are potentially more sub-cellular structures in this data (figure 4). However, the uncertainty is too great to support these phenotypes.

**Figure 4:**
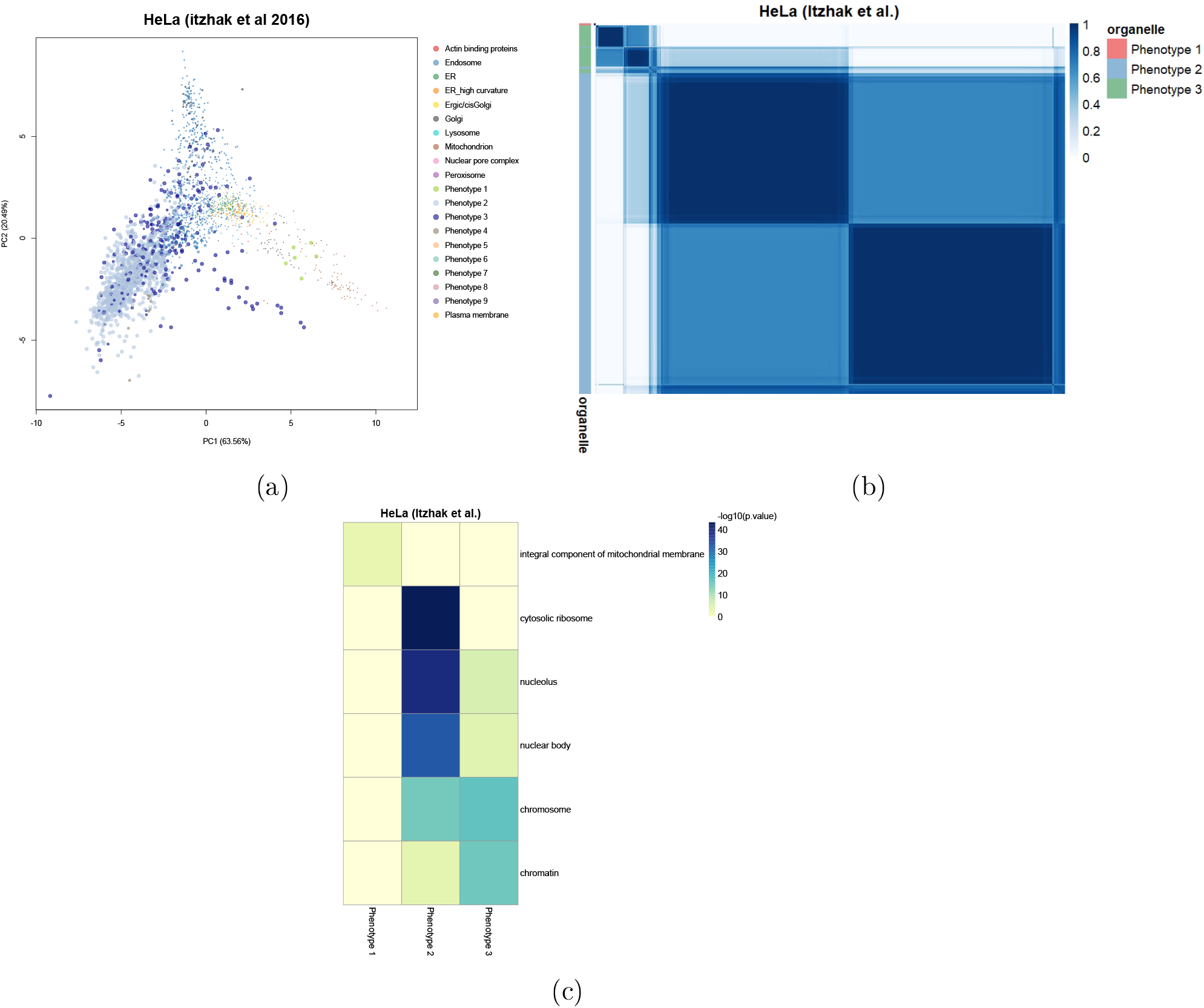
(a) PCA plots of the HeLa data. The pointers are scaled according to their discovery probability. (b) Heatmaps of the HeLa Itzhak data. Only the proteins with discovery probability greater than 0.99 and outlier probability less than 0.95 are shown. The heatmaps demonstrate the uncertainty in the clustering structure present in the data. (c) Tile plot of phenotypes against GO CC terms where the colour intensity is the −log_10_ of the p-value.

#### 3.3.2 Mouse primary neurons and HeLa cells (Hirst et. al 2018)

Application of Novelty TAGM to mouse primary neuron data (Itzhak *et al.*, 2017) and another HeLa dataset (Hirst *et al.*, 2018) yields further annotations; such as, *ribosomal, cytosolic* and *extracellular* annotations (see supplement 7.3).

## 4 Conparison between Novelty TAGM and *phenoDisco*

Next, we compare an already available novelty detection algorithm, *phenoDisco*, with Novelty TAGM. Despite both methods performing novelty detection, the algorithms are quite distinct. The first major difference is that Novelty TAGM is a Bayesian method that performs uncertainty quantification. Novelty TAGM quantifies the uncertainty in both the number of newly identified phenotypes and whether individual proteins should belong to a new phenotype. On the other hand, *phenoDisco* uses the *Bayesian Injormation Criterion* (BIC) to select just a single clustering, without taking into account the uncertainty in the number of phenotypes, and does not provide an estimate of individual protein-to-phenotype allocation uncertainty. Another difference is the input to both methods; Novelty TAGM uses the data directly, whereas *phenoDisco* takes the top principal components (by default, the first two) as input. *phenoDisco* also requires an additional parameter - the minimum group size. This parameter can be challenging to specify, since there is a trade-off between identifying functionally relevant phenotypes of different sizes and picking up small spurious protein clusters. Furthermore, *phenoDisco* struggles to scale to many of the datasets presented in this manuscript, because it requires iteratively refitting models and building of an outlier test statistic.

To demonstrate the differences between the two approaches, we apply *phenoDisco* and Novelty TAGM to the HEK-293 spatial proteomics dataset interrogated by Breckels *et al.* (2013). The PCA plots in figure 5 reveal broad similarities in the location of the discovered phenotypes. Novelty TAGM provides more information than *phenoDisco*; for example, we can scale the pointer size to the discovery probability. We note that both methods reveal 8 putative phenotypes in the data. Figure 5 (panels d and e) reveals the distribution of proteins across these phenotypes. We conclude that both approaches are able to discover small and large clusters, with both methods identifying phenotypes with a few proteins, but also phenotypes with greater than 100 proteins. Figure 5 (panel f) shows that both methods find the same number of phenotypes; however, not all of these phenotypes are functionally enriched. For *phenoDisco*, four of the phenotypes had at least 1 significant Gene Ontology term, whereas this was true for five of the Novelty TAGM phenotypes. Figure 5 (panel g) characterises the protein overlap between the two approaches. We see that both methods are in broad agreement, with most of the disagreement attributed to cases where one method assigns a protein as unknown whilst the other allocates to it a phenotype or organelle. For example, Novelty TAGM associates *phenoDisco* phenotype 3, which is a lysosome-enriched phenotype, with the plasma membrane (albeit with low probability). On the other hand, Novelty TAGM phenotypes 2 and 3, enriched for chromatin and ribosome respectively, are associated with the mitochondria by *phenoDisco*. This demonstrates the ability of Novelty TAGM to derive more biologically meaningful phenotypes.

**Figure 5:**
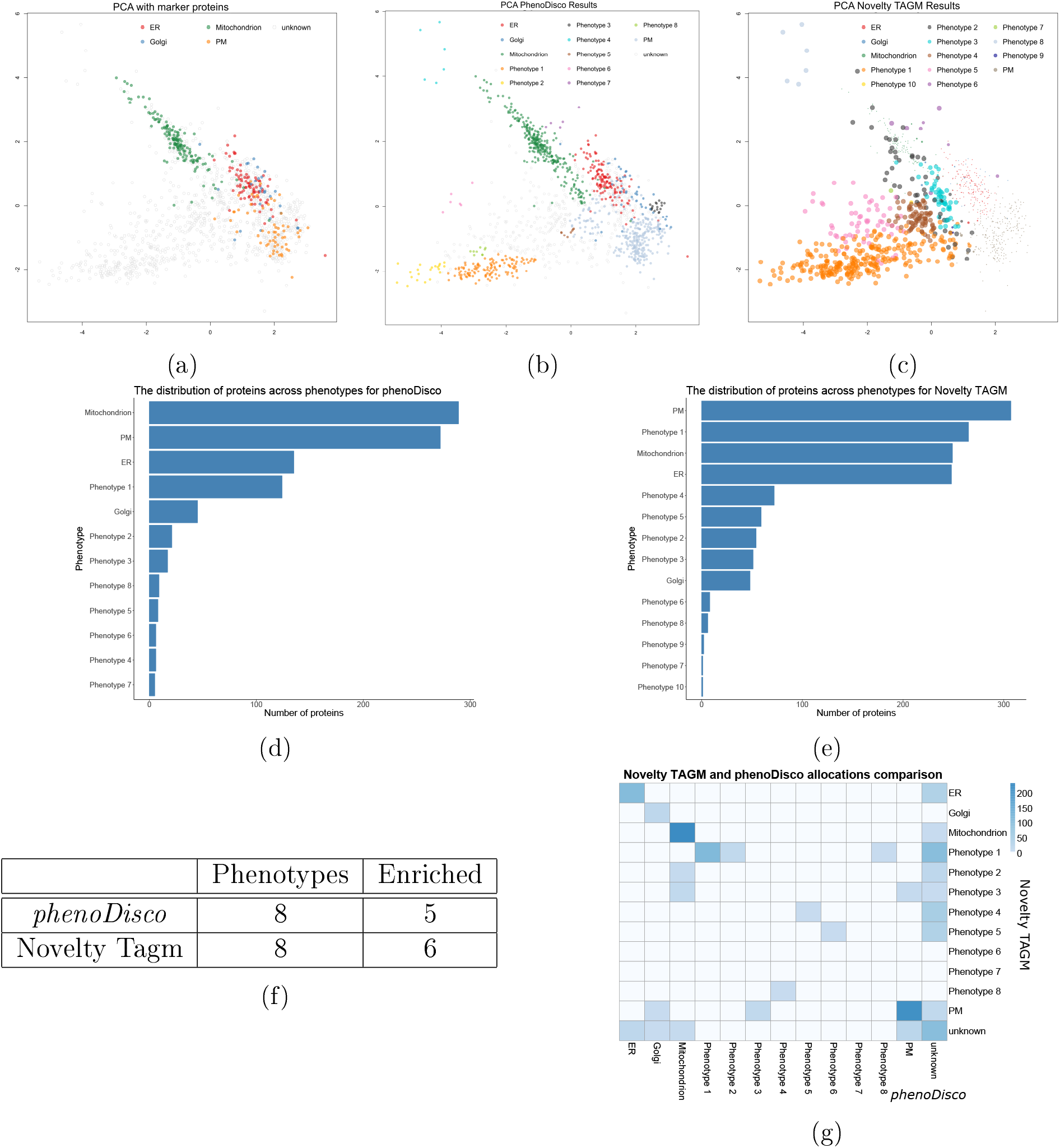
(a) PCA plot showing marker proteins for the HEK-293 dataset. (b) PCA plot with phenotypes identified by *phenoDisco*. (c) PCA plot with phenotypes identified by Novelty TAGM with pointer size scaled to discovery probability. (d, e) Barplots showing the number of proteins allocated to different phenotypes by *phenoDisco* and Novelty TAGM respectively. (f) A table demonstrating the number of phenotypes with functional enrichment for both methods and the number of phenotypes discovered. (g) A heatmap showing the overlap between *phenoDisco* and Novelty TAGM allocations.

## 5 Improved annotation allows exploration of endosonal processes

Given the information that the U-2 OS *hyper*LOPIT dataset resolves an endosomal cluster not previously explored, we perform a re-analysis of this dataset focusing on the endosomes. We curate a set of marker proteins for the endosomes and add these annotations to the U-2 OS *hyper*LOPIT dataset. After which, we apply our Bayesian generative classifier TAGM to the data with this additional annotation (Crook *et al.*, 2018). Protein allocations to each sub-cellular niche are visualised in the PCA plot of figure 6 (panel a). Figure 6 (panel c) demonstrates the increased number of proteins that can be characterised by improved annotation of the U-2 OS cell dataset. Furthermore, we examine 7 (of 240) proteins with uncertain endosomal localisation, which can be visualised in each of the violin plots in figure 6 (panel d).

**Figure 6:**
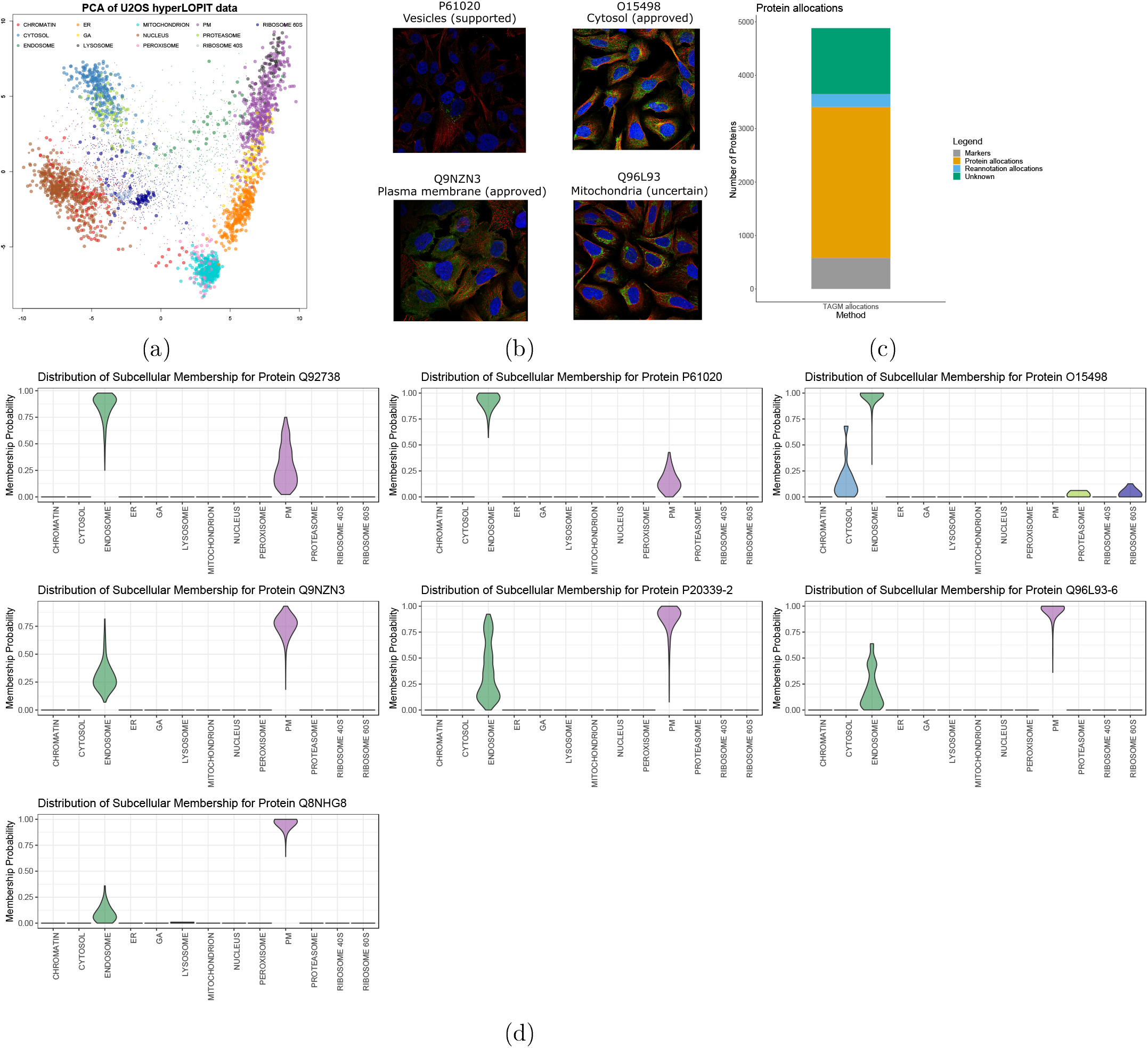
(a) PCA of U-2 OS *hyper*LOPIT data with pointer scaled to localisation probability and outliers shrunk. Points are coloured according to their most probable organelle. (b) Immunofluorescence images and sub-cellular localisation annotation taken from the HPA database (https://www.proteinatlas.org/humanproteome/cell) for the proteins with UniProt accessions P61020 (Rab5b), O15498 (Ykt6), Q9NZN3 (EHD3), and Q96L93 (KIF16B). The nucleus is stained in blue; microtubules in red, and the antibody staining targeting the protein in green. (c) A barplot representing the number of proteins allocated before and after re-annotation of the endosomal class. (d) Violin plots of full probability distribution of proteins to organelles, where each violin plot is for a single protein.

All 7 proteins with uncertain assignment to our new endosome cluster are known to function in endosome dynamics. Rab5a and Rab5b (P20339; P61020) are isoforms of Rab5, a small GTPase which is considered a master organiser of the endocytic system, regulating clathrin-mediated endocytosis and early endosome dynamics (Simonsen *et al.*, 1998; Woodman, 2000; Zerial and McBride, 2001; Rink *et al.*, 2005; Mendoza *et al.*, 2013; Chen *et al.*, 2014; Gautreau *et al.*, 2014; Law *et al.*, 2017). RN-tre (Q92738) is a GTPase-activating protein which controls the activity of several Rab GTPases, including Rab5, and is therefore a key player in the organisation and dynamics of the endocytic pathway (Lanzetti *et al.*, 2000; Gautreau *et al.*, 2014). KIF16B (Q96L93) is a plus end-directed molecular motor which regulates early endosome motility along microtubules. It is required for the establishment of the steady-state sub-cellular distribution of early endosomes, as well as the balance between PM recycling and lysosome degradation of signal transducing cell surface receptors including EGFR and TfR (Hoepfner *et al.*, 2005; Carlucci *et al.*, 2010). Notably, it has been demonstrated that KIF16B co-localises with the small GTPase Rab5, whose isoforms Rab5a and Rab5b we also identified as potentially localised to the endosome and PM in this dataset. ZNRF2 (Q8NHG8) is an E3 ubiquitin ligase which has been shown to regulate mTOR signalling as well as lysosomal acidity and homeostasis in mouse and human cells and has been detected at the endosomes, lysosomes, Golgi apparatus and PM according to the literature (Araki and Milbrandt, 2003; Hoxhaj *et al.*, 2016). Ykt6 (O15498) is a SNARE (soluble N-ethylmaleimide-sensitive factor attachment protein receptor) protein that regulates a wide variety of intracellular trafficking and membrane tethering and fusion processes. The membrane-associated form of Ykt6 has been detected at the PM, ER, Golgi apparatus, endosomes, lysosomes, vacuoles (in yeast), and autophagosomes as part of various SNARE complexes (Dilcher *et al.*, 2001; Tai *et al.*, 2004; Fukasawa *et al.*, 2004; Meiringer *et al.*, 2008; Takats *et al.*, 2018; Matsui *et al.*, 2018; Linnemannstöns *et al.*, 2018; Yong and Tang, 2019). In line with this, our results show a mixed sub-cellular distribution for Ykt6 with potential localisation to the endosome and cytosol (figure 6, panel d). EHD3 (Q9NZN3) is an important regulator of endocytic trafficking and recycling, which promotes the biogenesis and stabilisation of tubular recycling endosomes by inducing early endosome membrane bending and tubulation (Bahl *et al.*, 2016; Henmi *et al.*, 2016). We observe a mixed steady-state potential localisation to the endosome and PM for EHD3 (figure 6, panel d). This is in agreement with EHD3’s role in recycling endosome-to-PM transport (Naslavsky *et al.*, 2006, 2009; George *et al.*, 2007; Cabasso *et al.*, 2015; Henmi *et al.*, 2016).

Of these 7 proteins with uncertain endosome assignment, only 4 have localisations annotated in HPA (figure 6 (b)). The HPA assigns Rab5b to the vesicles which, in this context, include the endosomes, lysosomes, peroxisomes and lipid droplets. Therefore, a more precise annotation is available using Novelty TAGM. Ykt6 is localised to the cytosol, in support of our observations. EHD3 has approved localisation to the plasma membrane, again in agreement with our assignments. KIF16B is assigned to the mitochondrion, which contradicts our findings as well as previously published literature on the localisation and biological role of this protein. We speculate that this disagreement arises from the uncertainty associated with the specificity of the chosen antibody (Thul *et al.*, 2017). Thus, Novelty TAGM enables sub-cellular fractionation-based methods to identify proteins in sub-cellular niches which can not be fully interrogated by immunocytochemistry.

## 6 Discussion

We have presented a semi-supervised Bayesian approach that simultaneously allows probabilistic allocation of proteins to organelles, detection of outlier proteins, as well as the discovery of novel sub-cellular structures. Our method unifies several approaches present in the literature, combining the ideas of supervised machine learning and unsupervised structure discovery. Formulating inference in a Bayesian framework allows for the quantification of uncertainty; in particular, the uncertainty in the number of newly discovered annotations.

Application of our method across 10 different spatial proteomic datasets acquired using diverse fractionation and MS data acquisition protocols and displaying varying levels of resolution revealed additional annotation in every single dataset. Our analysis recovered the chromatin-associated protein phenotype and validated experimental design for chromatin enrichment in *hyper*LOPIT datasets. Our approach also revealed additional sub-cellular niches in the mESC *hyper*LOPIT and U-2 OS *hyper*LOPIT datasets.

Our method revealed resolution of 4 sub-nuclear compartments in the U-2 OS *hyper*LOPIT dataset, which were validated by Human Protein Atlas annotations. An additional endosome-enriched phenotype was uncovered and Novelty TAGM robustly identified an overlapping phenotype in U-2 OS LOPIT-DC data, providing strong evidence for endosomal resolution. Further biologically relevant annotations were uncovered in these, as well as other datasets. For example, a group of vesicle-associated proteins involved in transport from the ER to the early Golgi was identified in the yeast *hyper*LOPIT dataset; resolution of the ribosomal sub-units was identified in the fibroblast dataset, and separate nuclear, cytosolic and ribosomal annotations were identified in the DOM datasets.

A direct comparison with the state-of-the-art approach *phenoDisco* demonstrates clear differences between the approaches. Novelty TAGM, a fully Bayesian approach, quantifies uncertainty in both the number of newly discovered phenotypes and the individual protein-phenotype associations - *phenoDisco* provides no such information.

Improved annotation of the U-2 OS *hyper*LOPIT data allowed us to explore endosomal processes, which have not previously been considered with this dataset. We compare our results directly to immunofluorescence microscopy-based information from the HPA database and demonstrate the value of orthogonal spatial proteomics approaches to determine protein sub-cellular localisation. Our results provide insights on the sub-cellular localisation of proteins for which there is no information in the HPA Cell Atlas database.

During our analysis, we observed that the posterior similarity matrices have potential sub-clustering structures. Many known organelles and sub-cellular niches have sub-compartmentalisation, thus methodology to detect these sub-compartments is in preparation. Furthermore, we have observed that different experiments and different data modalities provide complementary results. Thus, integrative approaches to spatial proteomics analysis are also desired.

Our method is widely applicable within the field of spatial proteomics and builds upon state-of-the-art approaches. The computational algorithms presented here are disseminated as part of the Bioconductor project (Gentleman *et al.*, 2004; Huber *et al.*, 2013) building on MS-based data structures provided in Gatto and Lilley (2012) and are available as part of the pRoloc suite, with all data provided in pRolocdata (Gatto *et al.*, 2014b).

## 7 Appendix

## 7.1 mESC chromatin enrichment validation

For the mESC dataset, Novelty TAGM reveals 8 new putative phenotypes. Novelty TAGM recovers the masked annotations with phenotype 2 having the enriched terms associated with chromatin, such as *chromatin* and *chromosome* (*p <* 10^−80^). Phenotype 3 corresponds to a separate nuclear substructure with enrichment for the terms *nucleolus* (*p <* 10^−60^) and *nuclear body* (*p <* 10^−30^). Thus, in the mESC dataset Novelty TAGM confirms the chromatin enrichment preparation designed to separate chromatin and non-chromatin associated nuclear proteins (Mulvey *et al.*, 2017). In addition, phenotype 4 demonstrates enrichment for the ribosome annotation (*p <* 10^−35^). Phenotype 1 is enriched for *centrosome* and *microtubule* annotations (*p <* 10^−15^), though observing the PSM in figure 7 we can see there is much uncertainty in this phenotype. This uncertainty quantification can then be used as a basis for justifying additional expert annotation.

**Figure 7:**
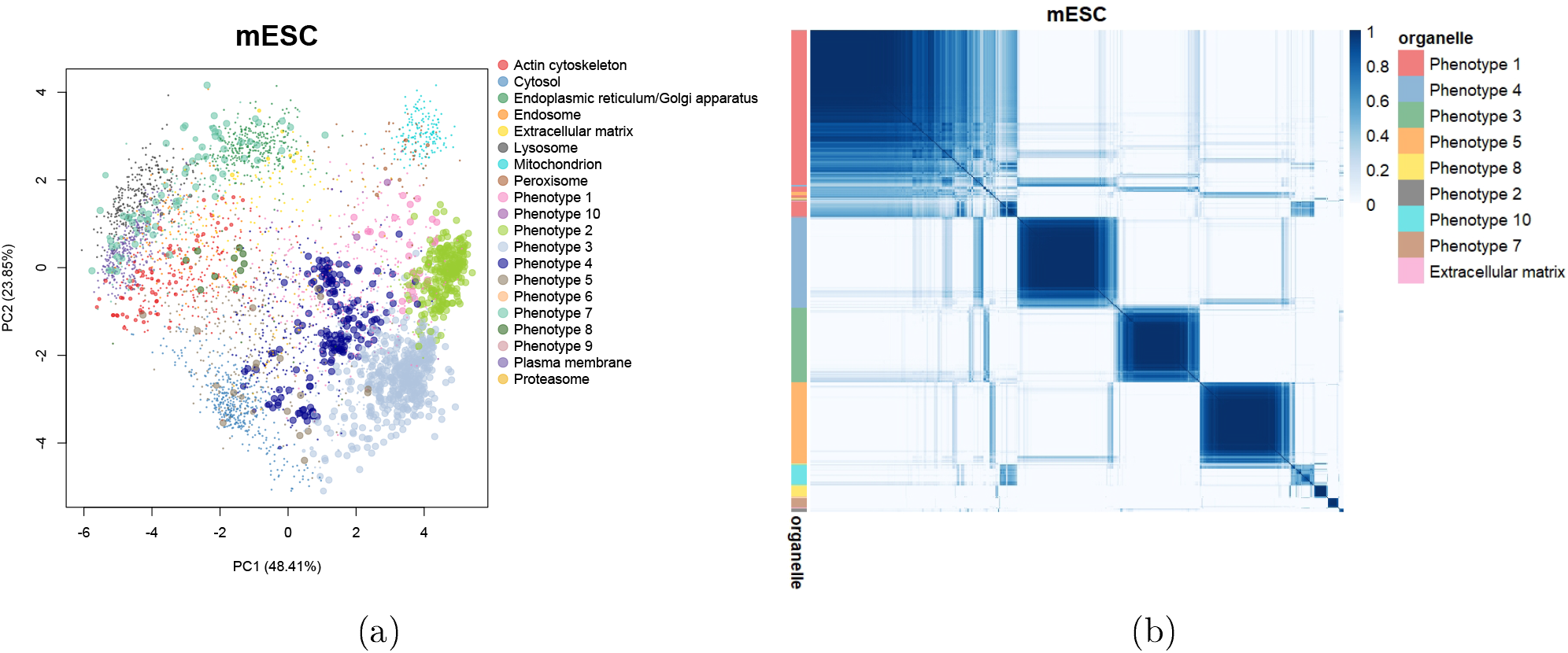
(a) PCA plot of the *hyper*LOPIT mESC dataset. Points are scaled according to the discovery probability. (b) Heatmaps of the posterior similarity matrix derived from mESC data demonstrating the uncertainty in the clustering structure of the data. We have only plotted the proteins which have greater than 0.99 probability of belonging to a new phenotype and probability of being an outlier less than 0.95 for the mESC dataset to reduce the number of visualised proteins.

## 7.2 Uncovering additional annotations in fibroblast cells

## 7.2.1 HCMV-infected fibroblast cells

We apply Novelty TAGM to the dataset corresponding to the HCMV-infected fibroblast cells 24 hours post infection (hpi) (Beltran *et al.*, 2016), and discover 9 putative additional phenotypes (demonstrated in figure 8). Phenotype 2 contains a singleton protein and phenotypes 4, 6, 7, 8 and 9 are not significantly enriched for any annotations. However, phenotype 3 is enriched for the *mitochondrial membrane* and *mitochondrial envelope* annotations (*p <* 10^−4^); this is an addition to the already annotated mitochondrial class, indicating sub-mitochondrial resolution. Phenotype 1 is a mixed ribosomal/nuclear cluster with enrichment for *nucleoplasm* (*p <* 10^−5^) and the *small ribosomal subunit* (*p <* 10^−4^), which is distinct from phenotype 5 which is enriched for the *large ribosomal subunit* (*p <* 10^−10^). This demonstrates unbiased separation of the two ribosomal subunits, which was overlooked in the original analysis (Beltran *et al.*, 2016).

## 7.2.2 Fibroblast cells without infection

Novelty TAGM reveals 7 putative phenotypes in the control fibroblast dataset (Beltran *et al.*, 2016). Phenotypes 2, 4, 5, 6 and 9 have no significantly enriched Gene Ontology terms (threshold *p* = 0.01). However, we observe that phenotype 3 is enriched with the *large ribosomal subunit* with significance at level *p <* 10^−7^. Phenotype 1 represents a mixed *peroxisome* (*p <* 10^−2^) and *mitochondrion* cluster (*p <* 10^−2^), an unsurprising result since these organelles possess similar biochemical properties and therefore similar profiles during density gradient centrifugation-based fractionation (Geladaki *et al.*, 2019; Dealtry and Rickwood, 1992). The differing number of confidently identified and biologically relevant phenotypes discovered between the two fibroblast datasets could be down to the differing levels of structure between the two datasets. Indeed, it is evident from figure 8 that we see differing levels of clustering structure in these datasets.

## 7.3 Additional organellar map datasets

## 7.3.1 Mouse primary neurons

The mouse primary neuron dataset reveals 10 phenotypes after we apply Novelty TAGM. However, 8 of these phenotypes have no enriched GO annotations. This is likely a manifestation of the dispersed nature of this dataset, where the variability is generated by technical artefacts rather than biological signal. Despite this, Novelty TAGM is able to detect two relevant phenotypes: the first phenotype is enriched for *nucleolus* (*p <* 0.01); the second for *chromosome* (*p <* 0.01). This suggests additional annotations for this dataset.

## 7.3.2 HeLa cells (Hirst et. al 2018)

The HeLa dataset of Hirst *et al.* (2018), which we refer to as HeLa Hirst, reveals 7 phenotypes with at least 1 protein with discovery probability greater than 0.95. However, three of these phenotypes represent singleton proteins. Phenotype 1 reveals mixed cytosol/ribosomal annotations with the terms *cytosolic ribosome* (*p <* 10^−30^) and *cytosolic part* (*p <* 10^−25^) significantly over-represented. There are no further phenotypes with enriched annotations (threshold *p* = 0.01), except phenotype 2 which represents a mixed extracellular structure/cytosol cluster. For example, the terms *extracellular organelle* (*p <* 10^−13^) and *cytosol* (*p <* 10^−10^) are over-represented.

**Figure 8:**
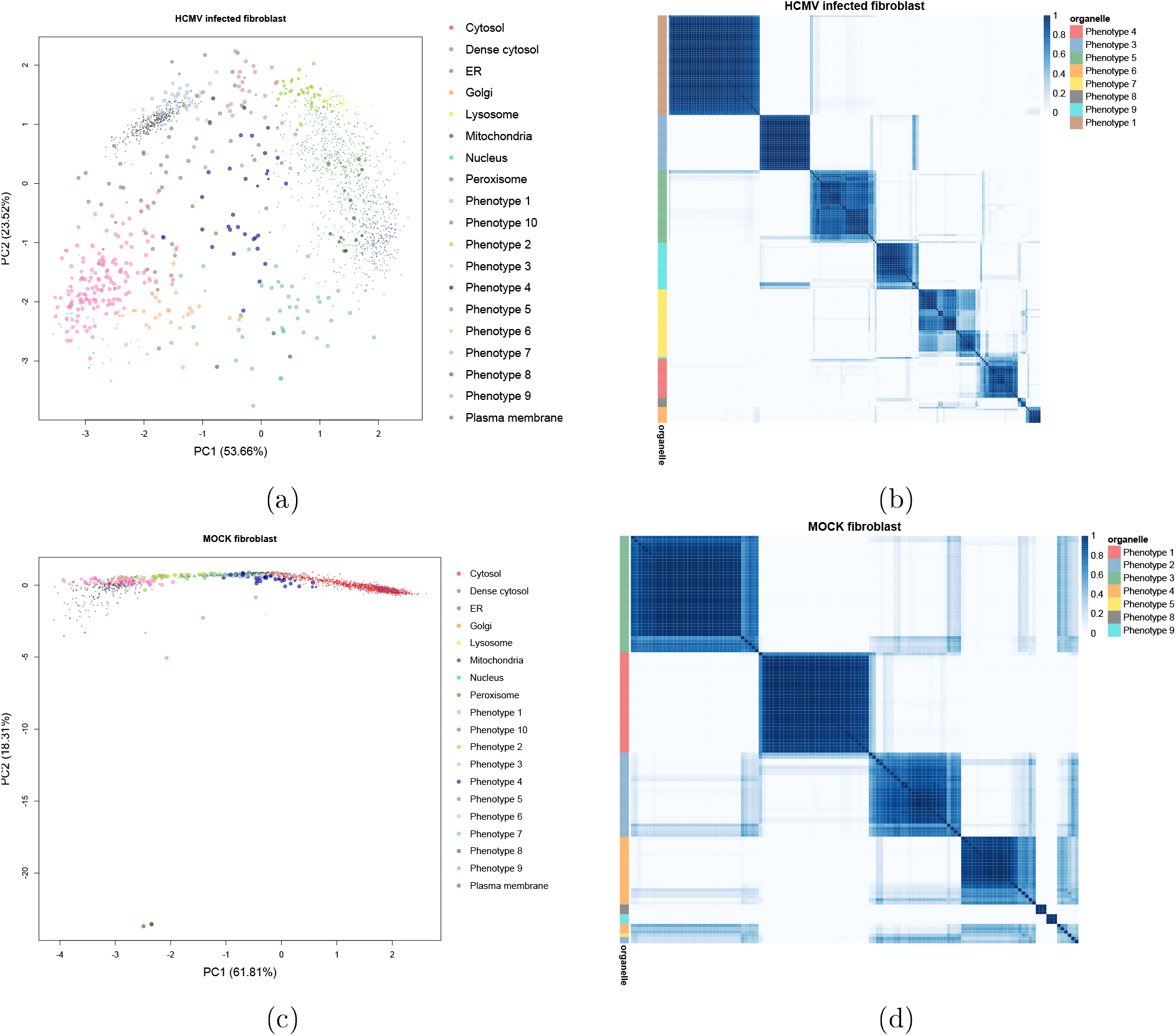
(a, c) PCA plots of the HCMV-infected fibroblast data 24 hpi and the mock fibroblast data 24 hpi. The points are coloured according to the organelle or proposed new phenotype and are scaled according to the discovery probability. (b, d) Heatmaps of the posterior similarity matrix derived from the infected fibroblast data and mock fibroblast data demonstrating the uncertainty in the clustering structure of the data. We have only plotted the proteins which have greater than 0.99 probability of belonging to a new phenotype and probability of being an outlier less than 0.95.

**Figure 9:**
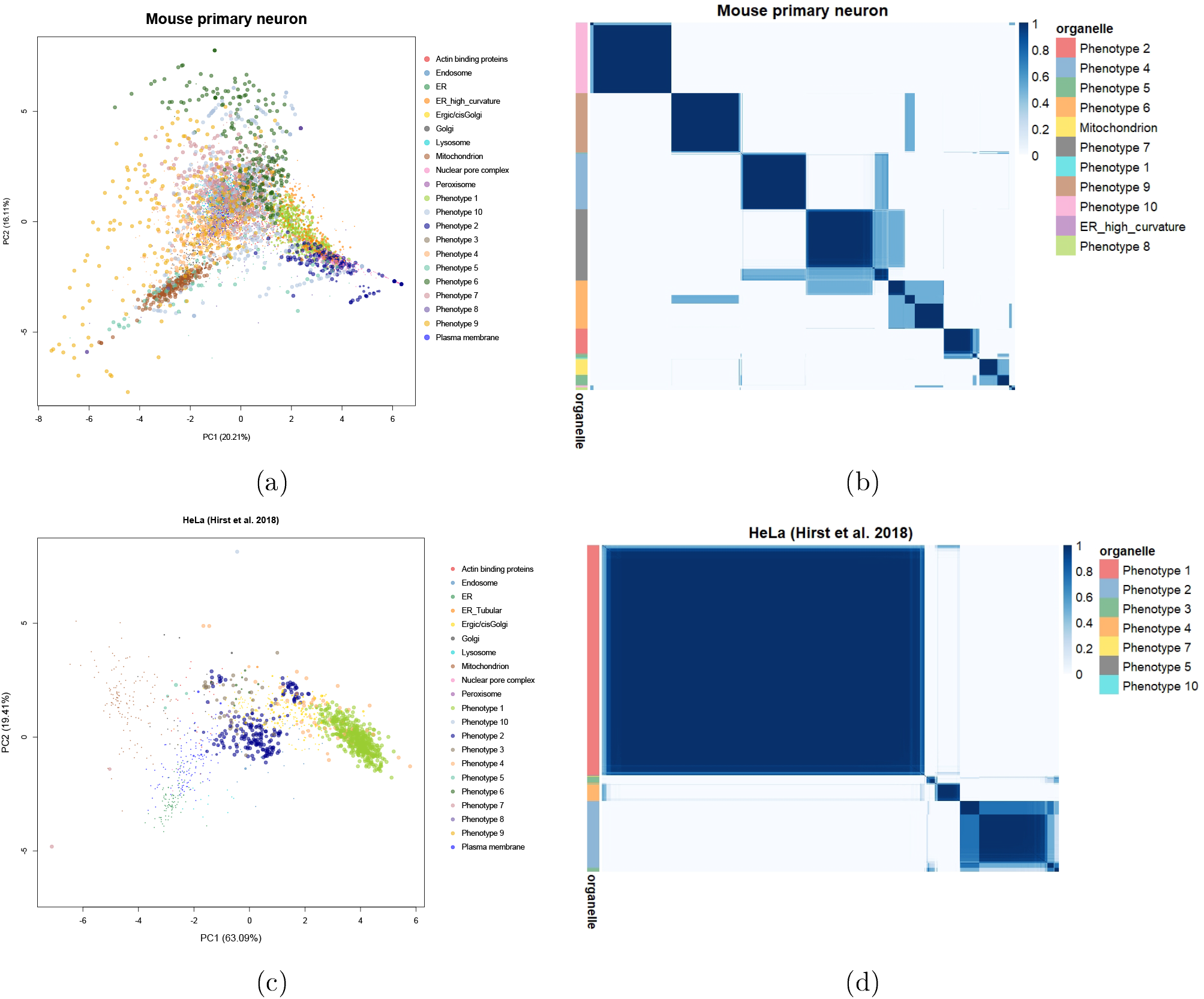
(a),(c) PCA plots of the mouse primary neuron data and HeLa Hirst data. The pointers are scaled according to their discovery probability. (b),(d) Heatmaps of the mouse neuron data and HeLa Hirst data. Only the proteins whose discovery probability is greater than 0.99 and outlier probability less than 0.95 (10^−2^ for the mouse primary neuron dataset to reduce the number of visualised proteins) are shown. The heatmaps demonstrate the uncertainty in the clustering structure present in the data.

## 7.4 Handling label switching

Bayesian inference in mixture models suffers from an identifiability issue known as *label switching* - a phenomenon where the allocation labels can flip between runs of the algorithm (Richardson and Green, 1997; Stephens, 2000). This occurs because of the symmetry of the likelihood function under permutations of these labels. We note that this only occurs in unsupervised or semi-supervised mixture models. This makes inference of the parameters in mixture models challenging. In our setting the labels for the known components do not switch, but for the new phenotypes label switching must occur. One standard approach to circumvent this issue is to form the so-called *posterior similarity matrix* (PSM) (Fritsch and Ickstadt, 2009). The PSM is an *N × N* matrix where the (*i, j*)^*th*^ entry is the posterior probability that protein *i* and protein *j* reside in the same component. More precisely, if we let *S* denote the PSM and *T* denote the number of Monte-Carlo iterations then

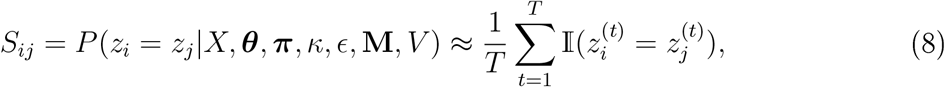

where 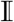 denotes the indicator function. The PSM is clearly invariant to label switching and so avoids the issues arising from the *label switching* problem.

## 7.5 Summarising posterior similarity matrices

To summarise the PSMs, we take the approach proposed by Fritsch and Ickstadt (2009). They proposed the adjusted Rand index (AR) (Rand, 1971; Hubert and Arabie, 1985), a measure of cluster similarity, as a utility function and then we wish to find the allocation vector 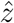 that maximises the expected adjusted Rand index with respect to the true clustering *z*. Formally, we write

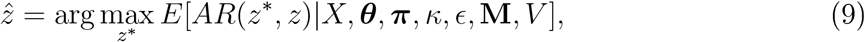

which is known as the Posterior Expected Adjusted Rand index (PEAR). One obvious pitfall is that this quantity depends on the unknown true clustering *z*. However, this can be approximated from the MCMC samples:

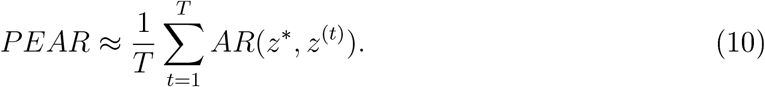

The space of all possible clustering over which to maximise is infeasibly large to explore. Thus we take an approach taken in Fritsch and Ickstadt (2009) to propose candidate clusterings over which to maximise. Using hierarchical clustering with distance 1 – *S_ij_*, the PEAR criterion is computed for clusterings at every level of the hierarchy. The optimal clustering 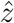 is the allocation vector which maximises the PEAR.

## 7.6 Details of MCMC

The MCMC algorithm used in Crook *et al.* (2018) is insufficient to handle inference of unknown phenotypes. As in Crook *et al.* (2018), a collapsed Gibbs sampler approach is used, but a number of modifications are made. Firstly, to accelerate convergence of the algorithm half the proteins are initial allocated randomly amongst the new phenotypes. Secondly, the parameters for the new phenotypes are proposed from the prior. Throughout the same default prior choices are used as in Crook *et al.* (2018).

## 7.7 Further details of endosomal proteins

For completeness, this appendix provides additional details and important literature on the proteins discussed in section 5.

First, P20339 (Rab5a) and P61020 (Rab5b) are two of the three isoforms of Rab5, a small GTPase which belongs to the Ras protein superfamily and is considered a master organiser of the endocytic system. Rab5a and Rab5b share a high level of amino acid sequence identity (approximately 85%) and are ubiquitously expressed in the mouse and human. Independently, these isoforms act as key regulators of clathrin-mediated endocytosis and early endosome dynamics by controlling the following processes *in vivo* and *in vitro*: (a) clathrin-coated vesicle formation at the cell surface; (b) endocytosed vesicle transport from the plasma membrane towards, and fusion with, early endosomes; (c) early endosome biogenesis and maintenance; (d) molecular motor-driven, microtubule-dependent early endosome motility along the endocytic route; (e) early endosome docking/tethering and homotypic fusion, and (f) Rab conversion and early-to-late endosome maturation (Simonsen *et al.*, 1998; Zerial and McBride, 2001; Rink *et al.*, 2005; Chen *et al.*, 2014; Gautreau *et al.*, 2014; Law *et al.*, 2017).

Rab5a and Rab5b play crucial roles in the internalisation and recycling/degradation of cell surface receptors such as EGFR (epidermal growth factor receptor), TfR (transferrin receptor) and several GPCRs (G-protein-coupled receptors) and integrins as well as peripheral plasma membrane-associated signalling molecules, thereby regulating important intracellular signal transduction pathways (Trischler *et al.*, 1999; Chen *et al.*, 2009; Bastin and Heximer, 2013; Liu *et al.*, 2015). We observe a mixed steady-state potential localisation between the endosome and PM for both Rab5a and Rab5b (figure 6, panel d). According to previously published information, both Rab5a and Rab5b are mainly localised to (and considered well-established constituents of) the early endosome compartment but have also been detected on the PM and clathrin-coated vesicles, in support of our results (Simonsen *et al.*, 1998; Woodman, 2000; Mendoza *et al.*, 2013). Moreover, according to the HPA Cell Atlas, Rab5b resides in the vesicles (which, in this context, include the endosomes, lysosomes, peroxisomes and lipid droplets). There is no information regarding the sub-cellular location of Rab5a in this database.

Second, Q92738 (RN-tre) is a GTPase-activating protein (GAP) which controls the activity of several Rab GTPases. RN-tre is a major Rab5 (see above) regulator and therefore a key player in the organisation and dynamics of the endocytic pathway (Lanzetti *et al.*, 2000; Gautreau *et al.*, 2014). This protein modulates the internalisation of and signal transduction mediated by cell surface receptors such as EGFR, TfR and *β*1 integrins (Lanzetti *et al.*, 2000; Martinu *et al.*, 2002; Palamidessi *et al.*, 2013; De Franceschi *et al.*, 2015). It also controls early endosome-to-Golgi retrograde transport and Golgi membrane organisation (Haas *et al.*, 2007). We observe a steady-state snapshot of the sub-cellular distribution of RN-tre with potential localisation to the endosome and PM (figure 6, panel d). In line with these results, RN-tre has been shown to reside in Rab5-positive early endosomes at steady state, but has also been detected at the PM and focal adhesions (Lanzetti *et al.*, 2000; Martinu *et al.*, 2002; Palamidessi *et al.*, 2013; Gautreau *et al.*, 2014; De Franceschi *et al.*, 2015). There is no information concerning the sub-cellular localisation of RN-tre in the HPA Cell Atlas database.

Third, Q96L93 (KIF16B) is a plus end-directed molecular motor which belongs to the kinesin-3 protein family. This kinesin regulates early endosome motility along microtubules and is required for the establishment of the steady-state sub-cellular distribution of early endosomes as well as the balance between PM recycling and lysosome degradation of signal transducing cell surface receptors including EGFR and TfR (Hoepfner *et al.*, 2005; Carlucci *et al.*, 2010). In neuronal cells, KIF16B plays an important role in the establishment of somatodendritic early endosome localisation and in the trafficking of AMPA and NGF receptors (Farkhondeh *et al.*, 2015). In epithelial cells, this protein controls the transcytosis of TfR from juxtanuclear recycling endosomes to apical recycling endosomes (Bay *et al.*, 2013). KIF16B is also involved in tubular endosome biogenesis and fission by regulating early endosome fusion (Skjeldal *et al.*, 2012). Lastly, this kinesin has been shown to mediate biosynthetic Golgi-to-endosome transport of FGFR (fibroblast growth factor receptor)-carrying vesicles and thereby control FGFR cell surface presentation and signalling during in vivo mouse embryogenesis (Ueno *et al.*, 2011). Our results indicate a mixed localisation to the endosome and PM for KIF16B (figure 6, panel d). In line with our observations, it has been reported that this protein is associated with early endosome membranes at steady state in mouse and human cells (Hoepfner *et al.*, 2005; Farkhondeh *et al.*, 2015). Additionally, it has been demonstrated that KIF16B co-localises with, and its spatial distribution and activity is regulated by, the small GTPase Rab5, whose isoforms Rab5a and Rab5b we also identified as potentially localised to the endosome and PM in the U-2 OS *hyper*LOPIT dataset (see above), on early endosomes (Hoepfner *et al.*, 2005; Skjeldal *et al.*, 2012). Taking the above into account, a mixed distribution between the endosome and PM is reflective of the molecular function of KIF16B. However, the HPA Cell Atlas database classifies KIF16B as a component of the mitochondria (figure 6, panel b), contradicting our findings as well as previously published information regarding the sub-cellular localisation and biological role of this protein. We speculate that this disagreement arises from the uncertainty associated with the specificity of the chosen antibody (Thul *et al.*, 2017). Indeed, the reliability of the mitochondrial annotation for KIF16B is classified as “uncertain” in this database.

Fourth, Q8NHG8 (ZNRF2) is an E3 ubiquitin ligase which has been shown to regulate mTOR signalling as well as lysosomal acidity and homeostasis in mouse and human cells (Hoxhaj *et al.*, 2016). This protein has been found to control the sub-cellular localisation and biological function of mTORC1, the V-ATPase and the Na+/K+-ATPase *α*1 (Hoxhaj *et al.*, 2012, 2016). ZNRF2 is membrane-associated but can be released into the cytosol upon phosphorylation by various kinases (Hoxhaj *et al.*, 2016). We observe a mixed steady-state distribution between the endosome and PM for this protein (figure 6, panel d). In support of this result, we find that ZNRF2 has been detected on the endosomes, lysosomes, Golgi apparatus and PM according to the literature (Araki and Milbrandt, 2003; Hoxhaj *et al.*, 2016). There is no information in regard to the sub-cellular location of ZNRF2 in the HPA Cell Atlas database.

Fifth, O15498 (Ykt6) is a SNARE (soluble N-ethylmaleimide-sensitive factor attachment protein receptor) protein which is conserved from yeast to humans. This protein regulates a wide variety of intracellular trafficking and membrane tethering and fusion processes including ER-to-Golgi vesicular transport, intra-Golgi traffic, retrograde Golgi-to-ER transport, retrograde endosome-to-TGN (trans-Golgi network) trafficking, homotypic fusion of ER membranes, Golgi-to-PM transport and exosome/secretory vesicle-PM fusion, Golgi-to-vacuole traffic (in yeast), homotypic vacuole fusion (in yeast), autophagosome formation and autophagosome-lysosome fusion (Dilcher *et al.*, 2001; Tai *et al.*, 2004; Takáts *et al.*, 2018; Matsui *et al.*, 2018; Linnemannstöns *et al.*, 2018; Yong and Tang, 2019). Ykt6 lacks a trans-membrane domain and is able to cycle between intracellular membranes and the cytosol in a palmitoylation- and farnesylation-dependent manner (Fukasawa *et al.*, 2004; Meiringer *et al.*, 2008). The membrane-associated form of Ykt6 has been detected on the PM, ER, Golgi apparatus, endosomes, lysosomes, vacuoles (in yeast), and autophagosomes as part of various SNARE complexes (Dilcher *et al.*, 2001; Tai *et al.*, 2004; Fukasawa *et al.*, 2004; Meiringer *et al.*, 2008; Takáts *et al.*, 2018; Matsui *et al.*, 2018; Linnemannstöns *et al.*, 2018; Yong and Tang, 2019). In line with this information, our results show a mixed sub-cellular distribution for Ykt6 with potential localisation to the endosome and cytosol (figure 6, panel d). The cytosolic localisation for Ykt6 is also supported by the HPA Cell Atlas annotation corresponding to this protein (figure 6, panel b), further reinforcing our findings.

Sixth, Q9NZN3 (EHD3) is another important regulator of endocytic trafficking and recycling. This protein promotes the biogenesis and stabilisation of tubular recycling endosomes by inducing early endosome membrane bending and tubulation (Bahl *et al.*, 2016; Henmi *et al.*, 2016). Additionally, EHD3 is essential for early endosome-to-recycling endosome transport, retrograde early endosome-to-Golgi traffic, Golgi apparatus morphology maintenance, and recycling endosome-to-PM transport (Naslavsky *et al.*, 2006, 2009; George *et al.*, 2007; Cabasso *et al.*, 2015; Henmi *et al.*, 2016). It plays an important role in the recycling of cell surface receptors and the biosynthetic transport of lysosome proteins (Naslavsky *et al.*, 2006, 2009; George *et al.*, 2007; Cabasso *et al.*, 2015). We observe a mixed steady-state potential localisation to the endosome and PM for EHD3 (figure 6, panel d). Our results are in agreement with previously published studies which have reported that EHD3 is resident in the early endosomes and recycling endosomes at steady state (Naslavsky *et al.*, 2006, 2009; George *et al.*, 2007; Cabasso *et al.*, 2015), and our PM localisation-related observation is supported by the HPA Cell Atlas-derived annotation for this protein (figure 6, panel b).

Our findings provide insights on the dynamic sub-cellular distribution of proteins which play important roles in development, physiology and disease. For example, Rab5/Rab5a has been identified as a master regulator of cancer cell migration, tumour invasion and dissemination programs *in vitro* and *in vivo*. It has been demonstrated that Rab5/Rab5a expression is dysregulated in many invasive human cancers, Rab5/Rab5a is overexpressed in metastatic foci compared to the matched primary tumours, and Rab5/Rab5a activity critically promotes the acquisition of invasive properties by poorly invasive tumour cell types (Torres *et al.*, 2010; Liu *et al.*, 2011, 2015; Mendoza *et al.*, 2013; Frittoli *et al.*, 2014; Díaz *et al.*, 2014; Saitoh *et al.*, 2017). Several publications have reported that elevated Rab5/Rab5a expression correlates with, and is predictive of, increased local invasiveness and metastatic potential, as well as poor patient prognosis in a variety of human cancer types (Yu *et al.*, 1999; Fukui *et al.*, 2007; Zhao *et al.*, 2010; Yang *et al.*, 2011; Mendoza *et al.*, 2013; Frittoli *et al.*, 2014; Díaz *et al.*, 2014; Igarashi *et al.*, 2017). Due to its established role in cancer progression and metastasis, Rab5/Rab5a is considered a fundamental cancer-associated protein and a potential diagnostic marker or therapeutic target (Frittoli *et al.*, 2014; Igarashi *et al.*, 2017). Recently, Rab5 was identified as a promising therapeutic target for colorectal cancer, as inhibition of Rab5 (and Rab7) activity led to elimination of colorectal cancer stem cells and disruption of colorectal cancer foci (Takeda *et al.*, 2019). Moreover, individual ablation of Rab5a but also Rab5b was shown to impair the invasion and dissemination ability of different cancer cell types (Frittoli *et al.*, 2014). In addition to its important role in cancer, there is some evidence suggesting that Rab5a might also be involved in the early pathogenesis of Alzheimer’s disease (Cataldo *et al.*, 1997, 2000; Rosenfeld *et al.*, 2001). Lastly, the Rab5 machinery has also been identified as an important factor in several bacterial, parasitic and viral infections. Bacterial pathogens such as *Mycobacterium tuberculosis, Listeria monocytogenes, Tropheryma whipplei* and *Salmonella typhimurium* (Madan *et al.*, 2008), as well as parasites such as *Leishmania donovani* have evolved specific subversion mechanisms with which they are able to control the intracellular distribution and/or activity of Rab5 and its effectors as a way to avoid neutralisation by the immune system or facilitate invasion (Verma *et al.*, 2017). *L. donovani* specifically controls the expression and function of the Rab5a isoform in this context (Verma *et al.*, 2017). Additionally, Rab5 was shown to participate in adenovirus endocytosis (Rauma *et al.*, 1999), both Rab5a and Rab5b were found to play functional roles in web formation and viral genome replication during HCV (hepatitis C virus) infection (Stone *et al.*, 2007), and Rab5a was identified as a crucial target of HBV (hepatitis B virus) during HBV-related hepatocellular carcinoma pathogenesis (Sheng *et al.*, 2014).

Apart from Rab5a and Rab5b, the other proteins also possess demonstrated roles in development and disease. RN-tre is overexpressed in a subset of aggressive basal-like breast cancers, where high levels of this protein prevent the endocytosis and recycling of EGFR, leading to Akt overstimulation. In turn, Akt activity stabilises the glucose transporter GLUT1 at the cell membrane, resulting in an increase in glycolysis and cancer cell proliferation. RN-tre has been proposed as a potential therapeutic target for these types of breast cancer (Avanzato *et al.*, 2018). This protein also plays a functional role in infection, as it was shown to regulate the uptake and intracellular trafficking of Shiga toxins (Fuchs *et al.*, 2007). Furthermore, it has been reported that KIF16B is essential for early post-implantation mouse embryo development, as Kif16b-knockout animals display peri-implantation embryonic lethality (Ueno *et al.*, 2011). In addition, recent studies have shown that ZNRF2 is overexpressed in human non-small cell lung cancer, osteosarcoma and papillary thyroid cancer, and that high levels of this protein are correlated with disease progression and poor patient prognosis in these cases (Zhang *et al.*, 2016; Xiao *et al.*, 2017; Cui *et al.*, 2019). Moreover, Ykt6 was found to be necessary for glycosome biogenesis and function in the kinetoplastid parasite *Trypanosoma brucei*, which causes African sleeping sickness, with Ykt6 ablation significantly reducing the viability of the parasite in both its pro-cyclic and bloodstream forms (Banerjee and Rachubinski, 2017). Finally, EHD3 has been identified as an essential factor for heart physiology (Curran *et al.*, 2014).

## 7.8 Summary of convergence diagnostics

We provide a summary of convergence diagnostics using parallel chains analysis (Gelman and Rubin, 1992). We compute the number of proteins allocated to the outlier component at each iteration of the Markov-chain and monitor this quantity for convergence. The 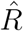 statistic between parallel chains in then computed and reported in the table below. A value of 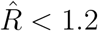 indicates convergence.

**Table 2:** A table reporting convergence diagnostics for MCMC analysis

| Convergence diagnostics for MCMC |  |  |  |
| --- | --- | --- | --- |
| Dataset | Protocol | $\hat{R}$ | Upper confidence Interval<br>$\hat{R}$ |
| mESC | *hyper*LOPIT | 1.03 | 1.15 |
| U-2 OS | *hyper*LOPIT | 1.00 | 1.00 |
| U-2 OS | LOPIT-DC | 1.02 | 1.06 |
| *S. cerevisiae* | *hyper*LOPIT | 1.00 | 1.01 |
| HCMV-infected fibroblast | Spatio-Temporal Proteomics | 1.01 | 1.02 |
| HCMV mock fibroblast | Spatio-Temporal Proteomics | 1.03 | 1.08 |
| HeLa (Itzhak *et al.*, 2016) | DOM | 1.07 | 1.21 |
| Mouse primary neurons | DOM | 1.04 | 1.13 |
| HeLa (Hirst *et al.*, 2018) | DOM | 1.02 | 1.06 |
| HEK-293 | LOPIT | 1.00 | 1.01 |

## 7.9 Prior specification and sensitivity

To complete the Bayesian specification, here we provide details of the priors on the model parameters. In the multivariate Gaussian components of the Novelty TAGM model, as with TAGM, a common and practical choice is the use of a normal-inverse-Wishart prior. That is,

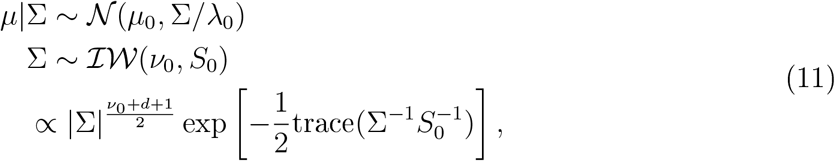

for each mixture component and where *d* is the dimension of the data. To complete this discussion, we need to specify the hyperparameters, *μ*_0_*, λ*_0_*, ν*_0_ and *S*_0_. We use diffusive priors that make minimal assumptions about the data, but they are set semi-empirically as to obtain the correct scale of the data. The hyperparameters are selected as follows

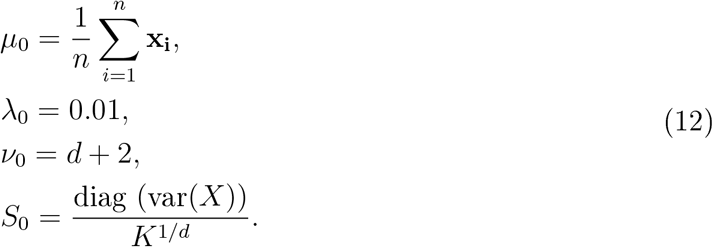

The hyperparameters are interpreted in the following ways. The prior mean, *μ*_0_, is the mean of the data. Then *λ*_0_ is viewed as the number of observations with data *μ*_0_ which are added to each component specific mean. This value is small to avoid strong prior influence. The marginal prior distribution (or prior predictive) for a cluster specific mean *μ* is given by a student’s *t*-distribution. This can be observed by recalling that the student’s *t*-distribution arises by marginalisation of the covariance from a normal distribution. Now, to ensure this *t*-distribution has finite covariance we require that *ν*_0_ *> d* + 1. Thus, the choice presented here is the smallest integer value of *ν*_0_ that ensures a finite covariance matrix. Hence, we have a well defined *t*-distribution with heavy tails. The empirically chosen scale matrix *S*_0_ is chosen to roughly partition the range of the data into *K* balls of equal size. Previous work has shown that these priors lead to good predictive performance (Crook *et al.*, 2018). For *π*, we take a conjugate symmetric Dirichlet prior with parameter *β*, so 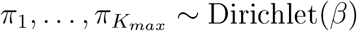. Note that to apply the principle of overfitted mixtures, we have to choose max_*j*_ *β*_j_ < *d*/2 (Rousseau and Mengersen, 2011), which is satisfied in all examples by setting *β_j_* = 1 for every *j*. Empirically Van Havre et al. (2013) have recommended smaller values of *β_j_ ≈ n*^−1^ to encourage stronger shrinkage.

## 7.9.1 Sensitivity to the choice of *β_j_*

To explore the sensitivity of our inferences to the specification of *β_j_*, we considered setting *β_j_* = 0.1, 0.01, as well as *β_j_ ≈ n*^−1^ for the mESC example, which in this case *n*^−1^ ≈ 0.0002. As before, we hid nucleus, chromatin and ribosome annotations and sought to use our model to rediscover them. As we now summarily describe, we found that our results can be sensitive to the choice of *β_j_* and hence it should be set carefully. For example when *β_j_* = 0.1, we were unable to detect a ribosomal phenotype. Furthermore, there was a joint nucleus and chromatin phenotype, phenotype 1, rather than two distinct phenotypes. *Chromosome* was enriched for this phenotype (*p <* 10^−100^), as well as nucleolus (*p <* 10^−60^). When *β_j_* = 0.01 the results were somewhat improved with a phenotype 1 enriched for *chromosome* (*p <* 10^−100^) but phenotype 3 was enriched for cytosolic ribosome (*p <* 10^−48^) and nucleolus (*p <* 10^−50^). Setting *β_j_* = 0.0002 provided the expected results with 3 distinct phenotypes for *chromatin* (phenotype 1) (*p <* 10^−100^), nucleolus (phenotype 4) (*p <* 10^−50^), and cytosolic ribsome (phenotype 3) (*p <* 10^−59^), successfully matching our test components. Hence, based on these results, we would recommend either *β_j_* = 1 or *β_j_≈ n*^−1^ depending on the desired amount of shrinkage.

## 7.10 Impact of reducing the proportion of labelled proteins

In all the examples we considered previously, the proportion of labelled proteins is roughly 20% of the total number of proteins. To assess the impact of the relative proportion of labelled and unlabelled proteins, we reconsidered our mESC example, where the goal was to detect ribosomal, nuclear and chromatin niches without annotation. In addition to masking these annotations as test components, we also masked, uniformly at random, an additional 10%, 20% and 50% of labelled proteins and assessed our ability to rediscover the ribosomal, nuclear and chromatin testing classes.

Briefly, we were able to rediscover two distinct phenotypes according to two nuclear clusters in all cases. When we masked 10% of the labels, the enrichments for the two nuclear phenotypes were chromosome (*p <* 10^−99^) and nucleolus (*p <* 10^−59^), the results were the same when we removed 20% and 50% of labels. However, only in the scenario were 20% of the labels were hidden did we find a ribosome enriched phenotype (*p <* 10^−30^). In the other cases, the ribosome clustered with the other large protein complex: the proteasome. This reflects the similar biochemical properties of these subcellular niches. Furthermore, removing annotations renders the proteasome profile less well defined, resulting in a more diffuse cluster. In practice, careful quality control would mitigate these scenarios (Gatto *et al.*, 2019). In applications where there are very few annotated niches and the analysis is close to the unsupervised setting, it may be valuable to increase *K_novelty_* above 10 - others have found *n/*2 to work well (Kirk *et al.*, 2012).

## 8 Acknowledgements

The authors would like to thank Tom Smith and Lisa M. Breckels of the Cambridge Centre for Proteomics for critical reading of the manuscript. This work was supported by the National Institute for Health Research [Cambridge Biomedical Research Centre at the Cambridge University Hospitals NHS Foundation Trust] [*]. *The views expressed are those of the authors and not necessarily those of the NHS, the NIHR or the Department of Health and Social Care.

## 9 Data Availability

All mass spectrometry data used in this manuscript is available in the *pRolocdata* Bioconductor package (Gatto *et al.*, 2014b). The Markov-chain Monte Carlo data can be found at Zenodo (Crook *et al.*, 2020a). Rmarkdown files to reproduce the figures can be found at the manuscript Github repository (Crook *et al.*, 2020b).

